# Conscious and unconscious eye contact at the limits of vision

**DOI:** 10.64898/2026.07.06.736643

**Authors:** Renzo C. Lanfranco, Arvid Guterstam, Axel Cleeremans

## Abstract

Eye contact is one of the most potent social signals, but it remains unclear how little visual input is sufficient to register direct gaze, and whether such registration requires conscious awareness. Here, we used a custom tachistoscope to present faces for only 1–5 ms, asking whether gaze direction can shape perception at the very limits of vision. Direct-gaze faces required less visual input to localise than averted-gaze faces and produced higher localisation sensitivity from 3 ms onwards. Crucially, this advantage emerged before participants could explicitly categorise gaze direction, and before perceptual awareness ratings showed metacognitive access to the information driving localisation, both of which appeared only at 4–5 ms. Information-theoretic analyses confirmed that localisation responses carried stimulus-location information before explicit reports and awareness ratings became informative. A second experiment showed that the earliest eye-contact advantage depended mainly on low spatial frequencies, indicating a role for coarse visual information. Finally, autistic traits were associated with a reduced localisation advantage for direct gaze. These findings show that eye contact can influence perceptual processing within just a few milliseconds, with less visual stimulation than is required for conscious access to gaze direction.

## INTRODUCTION

Eye contact is one of the most powerful and consequential signals in human social life[1, 2]. A direct gaze can initiate joint attention[3, 4], regulate affiliation and dominance[5], and rapidly influence affective and attentional processing[6–8]. Across development, faces that look directly at us preferentially capture attention and enhance social learning, suggesting that sensitivity to eye contact forms a foundational component of social cognition[9–11]. Yet a basic question remains unresolved: How much visual information is required for the brain to register that we are being looked at, and does this registration require conscious awareness?

In several domains of perception, the brain can extract information from visual input before that information becomes part of conscious experience[12]. This raises an intriguing possibility in the social domain: the visual system may detect that someone is looking at us before we consciously experience eye contact. Studies using interocular suppression techniques have suggested that faces with direct gaze gain access to awareness more readily than faces with averted gaze[13, 14]. However, such paradigms can neither determine whether direct gaze is genuinely detected from minimal visual input[15], nor whether their findings generalise beyond the specific suppression methods employed[16, 17]. The critical question, therefore, is whether the visual system registers eye contact before it reaches conscious awareness, and whether eye contact is prioritised over other facial cues during the emergence of conscious face perception.

One way to address this question is to estimate the minimal exposure thresholds required for different components of gaze processing[18–22]. Identifying such thresholds reveals whether eye contact processing and conscious awareness require the same minimal sensory input or whether they emerge at different levels of visual stimulation[15]. Here, we quantified the shortest display durations that evoked behavioural indices of gaze processing, thereby establishing the minimal visual input necessary to process eye contact and to consciously experience it. Using a high-precision tachistoscope[15, 23] capable of presenting images for exposures as brief as 0.002 ms[15], we isolated the earliest stages of visual processing without relying on masking or suppression techniques. If eye contact confers an early, awareness-independent processing advantage, then direct gaze should facilitate objective face detection at shorter exposure durations than those required for explicit gaze categorisation or metacognitive access.

To test this, we combined psychophysical and metacognitive measures. In Experiment 1, observers viewed intact faces and phase-scrambled control stimuli equated for lowlevel properties (Supplementary Note 1, Supplementary Fig. 1). We estimated perceptual thresholds and discrimination precision for multiple stimulus attributes, including the presence and location of a face, the direction of its gaze, and its emotional expression. Using signal-detection analyses[24, 25], we derived bias-free measures of sensitivity (*d*′) and, from psychometric functions, estimated minimal exposure thresholds and just-noticeable differences (JNDs). Participants also provided trial-by-trial ratings of their subjective visual experience[26, 27], allowing us to assess conscious access to perceptual evidence[28–30]. We further used information-theoretic analyses[31, 32] to quantify how much stimulus information was carried by objective localisation, explicit categorisation, and subjective awareness reports. This approach enabled us to determine whether objective sensitivity to eye contact emerged at shorter exposures than conscious access to gaze direction. We refer to such direct-gaze advantages over averted gaze as eye-contact effects[1]. In Experiment 2, we extended this design by applying spatial-frequency filtering to identify which visual channels primarily supported eye-contact processing across ultra-brief exposures. This allowed us to test whether coarse information often associated with subcortical pathways, or fine-grained detail often associated with cortical processing[1, 33–36] contributed differentially to eye-contact processing with ultra-brief exposures. We also asked whether the same spatial-frequency pattern was observed when performance was expressed as stimulus-response information rather than signal detection sensitivity. Finally, we assessed autistic traits in neurotypical adults to test whether individual differences predict eye contact processing at the limits of vision.

## RESULTS

### Determining the minimal exposure durations required to detect, categorise, and become aware of eye contact

In Experiment 1, we presented face images at ultra-brief exposure durations to determine how detection, categorisation, and conscious experience emerge as the amount of visual stimulation increases. Displays were presented for one of five equally spaced durations, from 1 to 5 ms, allowing us to test whether gaze direction and emotional expression modulated processing at the limits of vision (Fig. 1A,B). After each display, participants reported the location of the intact face, its gaze direction, and its emotional expression. They then rated their subjective visual experience of the intact face using a four-point perceptual awareness scale (PAS; Fig. 1C).

**Figure 1:**
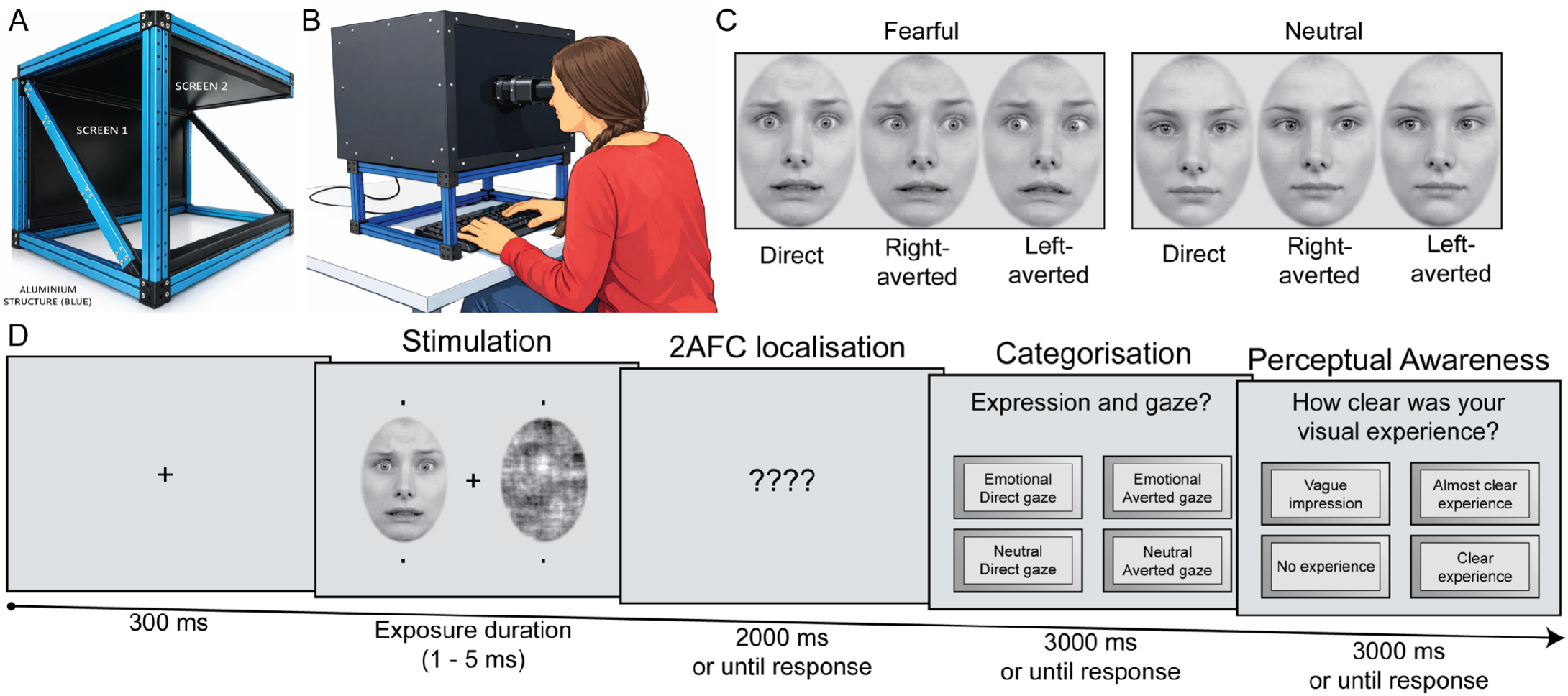
Schematic trial procedure of Experiment 1. **(A)** LCD tachistoscope used to present stimuli at ultra-brief exposure durations. The apparatus contains two LCD screens arranged orthogonally and combined optically through a semipermeable mirror. The horizontal screen presented target stimuli and its backlight was controlled by a dedicated microcontroller, enabling stimulus durations as brief as 0.002 ms. The vertical screen presented fixation displays, placeholders, response cues, and other non-target elements at regular presentation durations. **(B)** Participants viewed the superimposed display through a viewing aperture positioned 57 cm from the stimulus plane. **(C)** Face stimuli varied in emotional expression, fearful or neutral, and gaze direction, direct or averted. **(D)** On each trial, an intact face and its phase-scrambled counterpart were presented for a predefined exposure duration, 1 to 5 ms. Participants reported the location of the intact face, 2AFC, left or right, categorised its emotional expression, fearful or neutral, categorised its gaze direction, direct or averted, and rated their subjective visual experience of the intact face using a perceptual awareness scale, PAS. All face stimuli, including the intact face shown here, were taken from the Radboud Face Database (RaFD) and are presented as a stimulus example (see: https://rafd.nl/)

### Direct gaze reduces the sensory input required for face detection

We first asked whether gaze direction influenced the minimal visual input required to detect the location of an intact face. For each participant and condition, we fitted psychometric functions to localisation accuracy across the five exposure durations and estimated the exposure duration required to reach 75% correct performance (Fig. 2A). These threshold estimates revealed a significant effect of gaze direction (*F* _(1,29)_ = 4.26, *p* = 0.048, 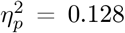), indicating that faces with direct gaze required less visual stimulation to be detected than faces with averted gaze. Emotional expression did not significantly affect detection thresholds, nor did it interact with gaze direction (see Supplementary Note 1 for additional details).

**Figure 2:**
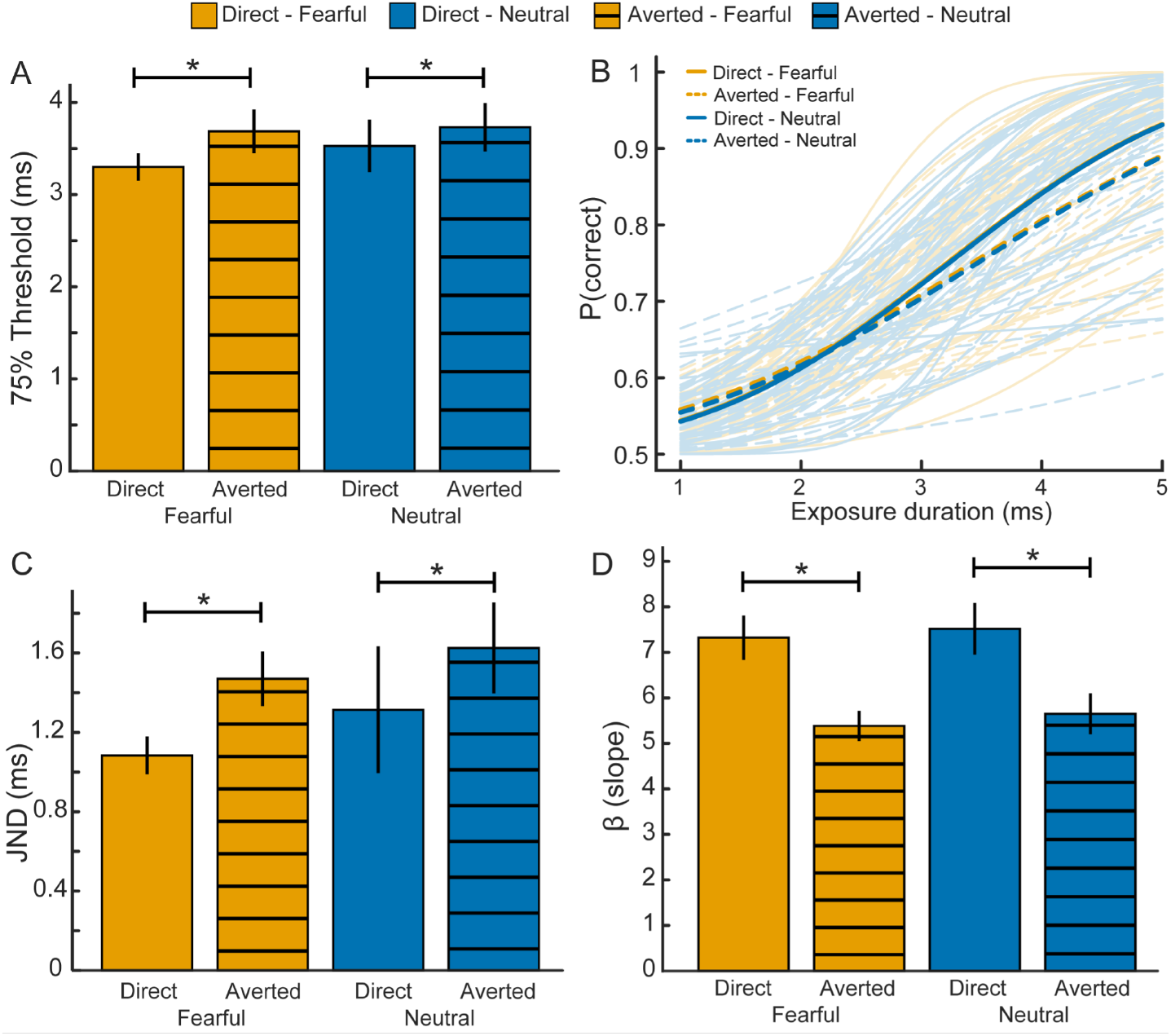
Psychometric evidence for a direct-gaze advantage in face detection at minimal visual stimulation. Exposure duration required to reach 75% correct localisation of the intact face, estimated from psychometric functions fitted separately for each gaze direction and emotional expression condition. Lower thresholds indicate that less visual stimulation was required for detection. Direct-gaze faces showed lower detection thresholds than averted-gaze faces, whereas emotional expression did not significantly affect thresholds. **(B)** Curve-level localisation accuracy across exposure durations. Accuracy increased as visual stimulation increased from 1 to 5 ms, and was higher for direct gaze than averted-gaze faces, confirming that gaze direction modulated detection performance across minimal exposures. **(C)** Just-noticeable differences estimated from the fitted psychometric functions. Smaller JNDs indicate greater discrimination precision. Gaze direction significantly affected JNDs, indicating greater precision for direct-gaze than averted-gaze faces. **(D)** Slope parameter estimates from the psychometric functions. Steeper slopes indicate a sharper increase in localisation performance as exposure duration increased. Slopes were significantly modulated by gaze direction, consistent with enhanced psychophysical performance for direct gaze. Error bars indicate *±*1 SEM. Individual lines in panel B indicate individual participants. Asterisks denote significant differences.

A curve-level analysis of localisation performance confirmed this pattern. Accuracy increased strongly as exposure duration increased (t(592) = 14.46, *p* < 0.001, *b* = 0.995, *SE* = 0.069, *OR* = 2.7; Fig. 2B), and was higher for direct-gaze than averted-gaze faces (t(592) = 3.32, *p* < 0.001, *b* = 0.142, *SE* = 0.043, *OR* = 1.15). Emotional expression did not predict localisation performance.

We next examined whether the gaze advantage also affected the precision of localisation performance. JNDs were significantly modulated by gaze direction (*F* _(1,29)_ = 5.24, *p* = 0.03, 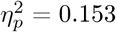; Fig. 2C), indicating greater localisation precision for direct-gaze than averted-gaze faces. Consistent with this precision effect, the slope parameter of the fitted psychometric functions was also strongly influenced by gaze direction (*F* _(1,29)_ = 37.49, *p* < 0.001, 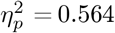 Fig. 2D). Emotional expression did not significantly affect either JNDs or slopes.

Thus, direct gaze reduced the visual input required for reliable localisation.

### The detection-discrimination-metacognitive access logic

Having established that direct gaze facilitated face localisation at minimal exposure durations, we then asked how this effect should be interpreted. A direct-gaze advantage in localisation does not, by itself, show that gaze information was processed without awareness, because better detection of one stimulus category over another could still depend on partial conscious access to the critical feature. A central challenge is therefore to determine whether the stimulus dimension that drives the detection advantage is also available for explicit report or subjective evaluation. We addressed this challenge by extending the detection-discrimination dissociation logic[37] with a metacognitive-access criterion (Fig. 3). In the standard detection-discrimination framework, evidence for unconscious processing is strongest when a stimulus dimension affects objective detection but participants cannot discriminate that dimension directly. However, chance-level discrimination does not fully rule out conscious access to the evidence driving performance. Participants may be unable to explicitly report the critical stimulus dimension, yet still have some subjective insight into whether their detection responses are likely to be correct. We therefore added a third requirement: subjective ratings should not reliably track the perceptual evidence supporting detection. Under this extended framework, the strongest evidence for unconscious eye-contact processing would be obtained if direct gaze improved face localisation while direct-versus-averted gaze categorisation remained unreliable and PAS ratings failed to distinguish correct from incorrect localisation responses. Together, these criteria tested whether eye contact influenced objective localisation before gaze information became available for explicit categorisation or metacognitive access.

**Figure 3:**
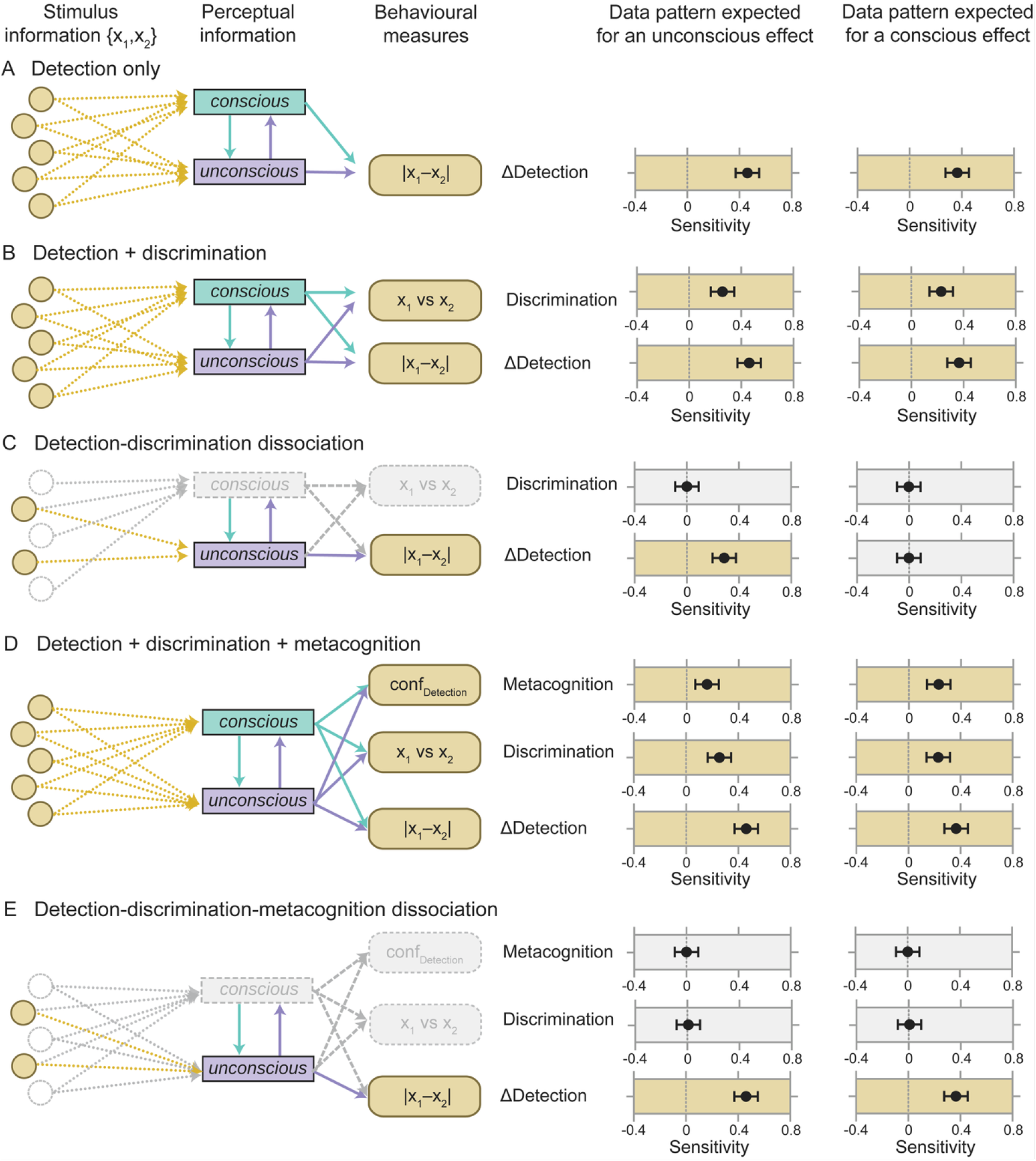
Extending the detection-discrimination dissociation logic to metacognitive access. Behavioural evidence can be used to distinguish conscious from unconscious contributions to detection effects. Stimulus information contains a critical dimension, {*x*_1_, *x*_2_}, such as upright versus inverted faces or direct versus averted gaze. This information may influence behaviour through conscious and/or unconscious perceptual processing. **(A)** In a standard detection paradigm, a positive detection difference between conditions, ΔDetection, indicates that {*x*_1_, *x*_2_} affects detection, but this alone does not reveal whether the effect depends on conscious or unconscious processing. **(B)** The detection-discrimination dissociation paradigm addresses this ambiguity by adding a direct discrimination measure for the critical dimension. If both ΔDetection and discrimination are above chance, conscious processing cannot be ruled out as the source of the detection effect. **(C)** If ΔDetection is above chance while discrimination of {*x*_1_, *x*_2_} remains at chance, the detection effect can be attributed to unconscious processing of the critical dimension. **(D)** We extend this logic by adding a metacognitive-access measure. In this extended framework, detection indexes whether the critical dimension influences behaviour, discrimination indexes whether that dimension is available for explicit report, and metacognitive access indexes whether subjective ratings track the perceptual evidence supporting detection performance. **(E)** The strongest evidence for unconscious processing is obtained when ΔDetection is above chance, but both discrimination and metacognitive access remain at chance. This pattern indicates that the critical stimulus dimension guides detection behaviour while remaining unavailable to explicit discrimination, and while subjective ratings fail to distinguish correct from incorrect detection responses. The right-hand columns show idealised data patterns expected for unconscious and conscious effects. Points and error bars indicate estimated effects and their uncertainty, with values around5zero reflecting chance-level performance or absent metacognitive sensitivity. Greyed elements indicate pathways or measures that are unavailable or non-informative in the corresponding idealised pattern. This figure extends schematics previously presented by Schmidt and Vorberg[38], and Stein and Peelen[37].

### Sensitivity to gaze direction and emotional expression

We used type-1 signal-detection analysis to estimate bias-free sensitivity to the relevant stimulus features. We computed *d*′ separately for localisation of the intact face, emotional expression categorisation, and gaze direction categorisation. See Supplementary Note 2 for more details.

Localisation sensitivity increased strongly with visual stimulation (*F* _(4,116)_ = 172.87, *p* < 0.001, 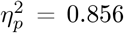), confirming that observers became progressively better at detecting the location of the intact face as exposure duration increased from 1 to 5 ms (Fig. 4A). Crucially, localisation sensitivity was also modulated by gaze direction (*F* _(1,29)_ = 25.37, *p* < 0.001, 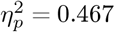), and this effect depended on exposure duration (*F* _(4,116)_ = 17.72, *p* < 0.001, 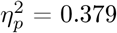). Follow-up comparisons showed no significant difference between direct and averted gaze at 1 or 2 ms, but significantly higher localisation sensitivity for direct-gaze faces at 3 ms (t_(29)_ = 4.04, Holm-corrected *p* = 0.001), 4 ms (t_(29)_ = 6.38, Holm-corrected *p* < 0.001), and 5 ms (t_(29)_ = 5.31, Holm-corrected *p* < 0.001). Emotional expression did not affect localisation sensitivity (*F* _(1,29)_ = 0.04, *p* = 0.838, 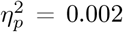), and did not interact with gaze direction or exposure duration (all *p* > 0.56). Localisation *d*′ was reliably above zero from 2 ms onwards in all gaze and expression conditions, but not at 1 ms. Thus, observers could detect the presence and location of an intact face with as little as 2 ms of visual stimulation, and direct gaze enhanced this sensitivity from 3 ms onwards.

**Figure 4:**
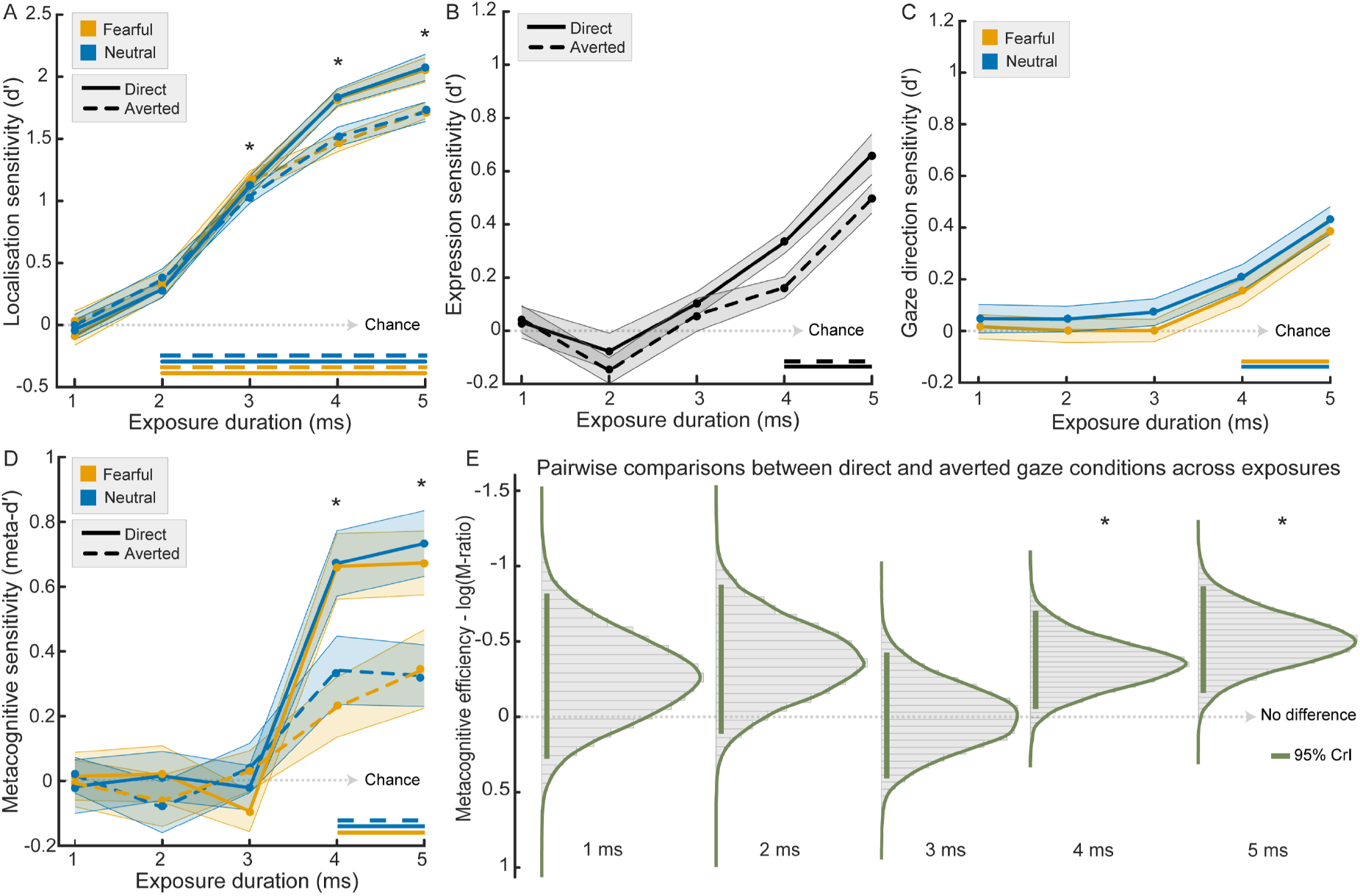
Type-1 and type-2 sensitivity across minimal exposure durations in Experiment 1. **(A)** Localisation sensitivity, indexed by type-1 *d*′, for detecting the position of the intact face. Sensitivity increased with exposure duration and was higher for direct gaze than averted-gaze faces from 3 ms onwards. **(B)** Expression sensitivity, indexed by type-1 *d*′, for categorising fearful versus neutral facial expressions. Expression sensitivity increased with exposure duration and became reliably above chance at longer exposures. **(C)** Gaze direction sensitivity, indexed by type-1 *d*′, for categorising direct versus averted gaze. Gaze categorisation improved with increasing visual stimulation and became reliable only at longer exposure durations. **(D)** Metacognitive sensitivity, estimated using maximum likelihood meta-*d*′, based on participants’ perceptual awareness ratings. This measure quantifies the extent to which subjective experience tracked objective perceptual evidence across exposure durations. Metacognitive sensitivity increased with exposure duration and was higher for direct gaze than averted-gaze faces from 4 ms onwards. **(E)** Metacognitive efficiency, estimated using Bayesian hierarchical M-ratio. Higher values indicate a closer correspondence between type-1 sensitivity and metacognitive access to that evidence. Posterior contrasts were assessed using 95% highest-density intervals, HDI. Contrasts whose confidence intervals included zero suggest no reliable difference between conditions. Metacognitive access to type-1 evidence was higher for direct-gaze faces from 4 ms onwards. Horizontal lines on top of the x-axis of Panels A–D indicate exposure durations with above-chance sensitivity, *p <* 0.05, one-sample *t*-tests against zero, Holm-corrected. Data are presented as mean *±*1 SEM, with shaded areas indicating SEM. Asterisks indicate statistically significant effects or planned comparisons after correction for multiple comparisons.

Expression categorisation also improved as visual stimulation increased (*F* _(4,116)_ = 36.99, *p* < 0.001, 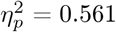). Expression sensitivity was modulated by gaze direction (*F* _(1,29)_ = 8.39, *p* = 0.007, 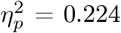), with higher sensitivity for direct than averted gaze faces (Fig. 4B). This effect did not interact with exposure duration (*F* _(4,116)_ = 1.29, *p* = 0.280, 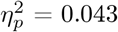). Against-zero tests indicated that expression sensitivity became reliable at 4 ms and 5 ms for both direct and averted gaze faces, but not at shorter exposure durations after correction.

Finally, gaze direction categorisation sensitivity increased with exposure duration (*F* _(4,116)_ = 22.80, *p* < 0.001, 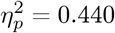 Fig. 4C). In contrast to localisation sensitivity, gaze categorisation *d*′ was not significantly modulated by emotional expression (*F* _(1,29)_ = 2.80, *p* = 0.105, 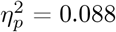), and there was no interaction between expression and exposure duration (*F* _(4,116)_ = 0.05, *p* = 0.993, 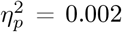). Gaze categorisation became reliably above chance only at 4 ms and 5 ms, for both fearful and neutral faces. Thus, direct gaze enhanced face localisation at exposure durations where explicit gaze direction categorisation was not yet reliable, indicating that eye contact influenced detection before observers could accurately categorise gaze direction. See Supplementary Fig. 2 for additional results.

### Dissociation between detection and conscious awareness

Next, we tested whether the perceptual evidence supporting face localisation was accompanied by conscious access to that evidence. To do so, we estimated metacognitive sensitivity using meta-*d*′, which quantifies how well subjective awareness tracks objective task performance[28, 39]. Meta-*d*′ was estimated from participants’ trial-by-trial perceptual awareness ratings. See Supplementary Note 3 for additional results. Whereas type-1 localisation sensitivity was already reliably above chance from 2 ms onwards, metacognitive sensitivity emerged later, indicating that observers could detect the intact face before their subjective experience reliably tracked the available perceptual evidence (Fig. 4D).

Meta-*d*′ increased significantly with visual stimulation (*F* _(4,108)_ = 25.26, *p* < 0.001, 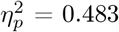). Metacognitive sensitivity was also strongly modulated by gaze direction (*F* _(1,27)_ = 18.24, *p* < 0.001, 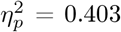), and this effect depended on exposure duration (*F* _(4,108)_ = 6.43, *p* < 0.001, 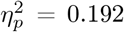). Follow-up comparisons showed no reliable difference between direct and averted gaze at 1 ms, 2 ms, or 3 ms (all Holm-corrected *p ≥*0.217). By contrast, meta-*d*′ was significantly higher for direct than averted gaze at 4 ms (t(29) = 4.31, Holm-corrected *p* < 0.001), and 5 ms (t(29) = 3.20, Holm-corrected *p* = 0.013). Emotional expression did not significantly affect meta-*d*′ (*F* _(1,27)_ = 0.41, *p* = 0.528, 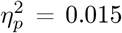), and did not interact with gaze direction or exposure duration (all *p* > 0.83).

Against-zero tests confirmed that metacognitive sensitivity was absent at the shortest exposure durations. Meta-*d*′ did not differ reliably from zero at 1, 2, or 3 ms in any condition after correction for multiple comparisons. For direct-gaze faces, meta-*d*′ became reliably positive at 4 ms and remained significant at 5 ms, for both fearful and neutral expressions. For averted-gaze faces, meta-*d*′ became reliable later and less consistently, reaching significance at 5 ms for fearful faces and at 4 and 5 ms for neutral faces. Thus, conscious access to face location emerged only at longer exposure durations than those required for type-1 localisation, and was stronger for faces making eye contact.

Meta-*d*′ indicates how much information about task performance is available to subjective awareness, but it is influenced by the amount of type-1 perceptual evidence available. We therefore also examined metacognitive efficiency, which asks whether subjective awareness tracks perceptual evidence more effectively for direct than averted gaze. We estimated M-ratio using a hierarchical metacognitive efficiency model, which quantifies how closely subjective awareness tracks type-1 perceptual sensitivity[29, 39] (Fig. 4E). The distribution of the posterior densities of M-ratio between direct and averted gaze did not differ at 1, 2, or 3 ms. However, M-ratio was significantly higher for direct-gaze than averted-gaze faces at 4 and 5 ms. Thus, once visual stimulation was sufficient to support reliable awareness, direct gaze was associated with more efficient mapping between perceptual evidence and subjective experience.

Together, these results reveal a dissociation between detection (assessed through type-1 *d*′) and conscious awareness at the limits of visual stimulation. Observers could localise intact faces from 2 ms onwards, and direct gaze enhanced localisation sensitivity from 3 ms onwards, but metacognitive sensitivity and metacognitive efficiency only became reliable from 4 ms onwards. Eye contact therefore influenced objective detection before subjective awareness reliably tracked the available perceptual evidence. See Supplementary Figs. 3 and 4 for additional results.

### Information-theoretic analyses confirm that eye contact increases localisation information

To complement the signal-detection analyses, we quantified the information carried by behavioural responses about the task-relevant stimulus variables[40, 41]. For each participant and for each exposure duration, we estimated bias-corrected mutual information between the relevant stimulus feature and the corresponding response. In the localisation task, this measured how much information participants’ responses carried about the actual location of the intact face, *I* (stimulus location; localisation response). In the expression and gaze tasks, we analogously estimated the information carried by categorisation responses about emotional expression and gaze direction. To relate objective localisation information to subjective experience, we also computed co-information between stimulus location, localisation response and PAS ratings, *CoI* (stimulus location; localisation response; PAS).

Localisation responses carried increasing information about the actual position of the intact face as exposure duration increased (Fig. 5A). Localisation information was not reliably above zero at 1 ms, became reliable in three of the four expression-by-gaze conditions at 2 ms, and was reliable in all four conditions from 3 ms onwards, reaching approximately 0.47–0.62 bits at 5 ms.

**Figure 5:**
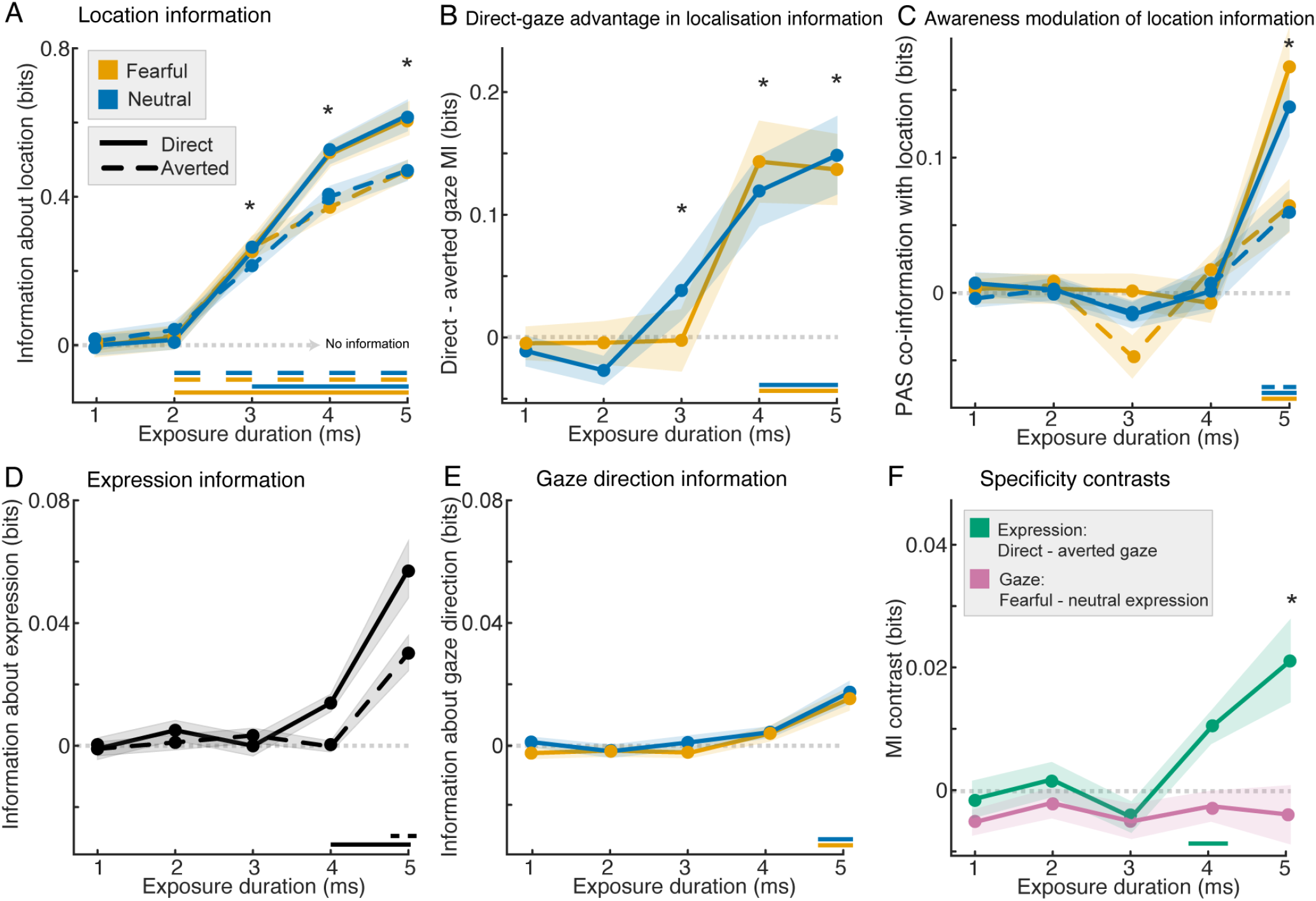
Information-theoretic evidence that eye contact increases localisation information. **(A)** Mutual information between the actual location of the intact face and participants’ localisation responses, *I*(stimulus location; localisation response), plotted separately by emotional expression and gaze direction. Localisation responses carried increasing information about stimulus location as exposure duration increased. **(B)** Direct-gaze advantage in localisation information, computed as the direct-minus-averted difference in localisation mutual information. Direct gaze increased localisation information at 4 ms and 5 ms for both fearful and neutral faces. **(C)** Co-information between stimulus location, localisation response, and PAS ratings, *CoI*(stimulus location; localisation response; PAS), shown separately for the four expression-by-gaze conditions, indexing the extent to which subjective awareness ratings overlapped with or modulated localisation information. Reliable positive co-information emerged at 5 ms for fearful-direct, neutral-direct, and neutral-averted faces, indicating that subjective awareness became coupled to localisation information at the longest exposure duration. **(D)** Mutual information between emotional expression and expression categorisation responses, *I*(expression shown; expression reported), plotted separately for direct- and averted-gaze faces. **(E)** Mutual information between gaze direction and gaze categorisation responses, *I*(gaze direction; gaze direction response), plotted separately for fearful and neutral faces. **(F)** Specificity contrasts for expression and gaze categorisation information. The expression contrast shows the direct-minus-averted difference in expression information; the gaze contrast shows the fearful-minus-neutral difference in gaze-direction information. Lines indicate group means and shaded areas indicate within-subject SEM. Dotted horizontal lines indicate zero information or zero contrast. Horizontal markers indicate exposure durations at which information estimates were significantly greater than zero after Holm correction. Asterisks indicate significant planned direct-minus-averted contrasts after Holm correction. For PAS co-information, the direct-minus-averted contrast at 5 ms indicates stronger coupling between subjective awareness ratings and localisation information for direct than averted gaze.

Crucially, direct gaze increased localisation information at the longer exposure durations. Planned direct-minus-averted contrasts were not reliable at 1, 2 or 3 ms after correction (Fig. 5B). However, localisation responses carried more information about stimulus location for direct-gaze than averted-gaze faces at both 4 and 5 ms. This was observed for fearful faces at 4 (t(29) = 3.9, Holm-corrected *p* = 0.0048, dz = 0.71) and 5 ms (t(29) = 4.19, Holm-corrected *p* = 0.002, dz = 0.77), and for neutral faces at 4 (t(29) = 3.89, Holm-corrected *p* = 0.0048, dz = 0.71) and 5 ms (t(29) = 4.37, Holm-corrected p = 0.0014, dz = 0.8). These results provide an information-theoretic counterpart to the signal detection findings: eye contact increased the amount of stimulus-location information available in objective localisation responses.

We next tested whether subjective awareness was coupled to localisation information (Fig. 5C). Co-information between stimulus location, localisation response and PAS ratings did not differ reliably from zero at 1–4 ms in any condition after Holm correction. At 5 ms, however, PAS co-information became reliably positive for fearful-direct faces (t(29) = 4.94, Holm-corrected *p* = 0.001, dz = 0.9), neutral-direct faces (t(29) = 5.55, Holm-corrected *p* < 0.001, dz = 1.01), and neutral-averted faces (t(29) = 3.98, Holm-corrected *p* = 0.016, dz = 0.73). The corresponding effect for fearful-averted faces was positive but did not survive correction. Thus, subjective visibility ratings became coupled to localisation information primarily at the longest exposure duration, indicating that awareness overlapped with, or modulated, the information shared between actual stimulus location and localisation responses once visual stimulation was sufficiently strong.

Expression and gaze categorisation showed more restricted information profiles (Fig. 5D,E). Expression information was not reliably above zero at 1–3 ms, became reliable for direct-gaze faces at 4 and 5 ms, and for averted-gaze faces only at 5 ms. Gaze-direction information was weaker still, becoming reliable only at 5 ms for both fearful and neutral faces. Specificity contrasts likewise indicated that the strongest information-theoretic modulation concerned localisation rather than explicit categorisation (Fig. 5F). Overall, direct gaze enhanced objective localisation information at the limits of vision, whereas explicit categorisation and subjective awareness emerged later. Full information-theoretic statistics are reported in Supplementary Note 4.

### Distinct contributions of coarse and fine spatial frequency information

In Experiment 2, we used type-1 signal-detection analysis to determine which spatial frequencies supported face localisation, the eye-contact effect, expression categorisation and gaze-direction categorisation under ultra-brief exposures. The design was similar to Experiment 1, but included only two exposure durations, 3 and 5 ms, and added spatial frequency as a within-participant factor, with low-spatial-frequency (LSF) and high-spatial-frequency (HSF) face images.

Localisation sensitivity increased strongly with exposure duration and was markedly higher for LSF than HSF faces (Exposure: *F* _(1,29)_ = 538.82, *p* < 0.001, 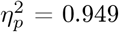 Spatial frequency: *F* _(1,29)_ = 232.64, *p* < 0.001, 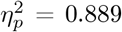 Fig. 6A,B). Localisation sensitivity was also higher for direct-gaze than averted-gaze faces (*F* _(1,29)_ = 57.82, *p* < 0.001, 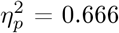), and this gaze effect depended on spatial frequency and exposure duration (Gaze direction *×* Spatial frequency: *F* _(1,29)_ = 5.06, *p* = 0.032, 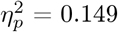). Condition-wise tests confirmed that localisation sensitivity was reliable for all LSF conditions at both exposure durations, whereas HSF localisation was weaker at 3 ms and became more consistently reliable at 5 ms (see Supplementary Fig. 5 for additional results). Thus, face localisation under ultra-brief exposure depended strongly on coarse spatial information.

**Figure 6:**
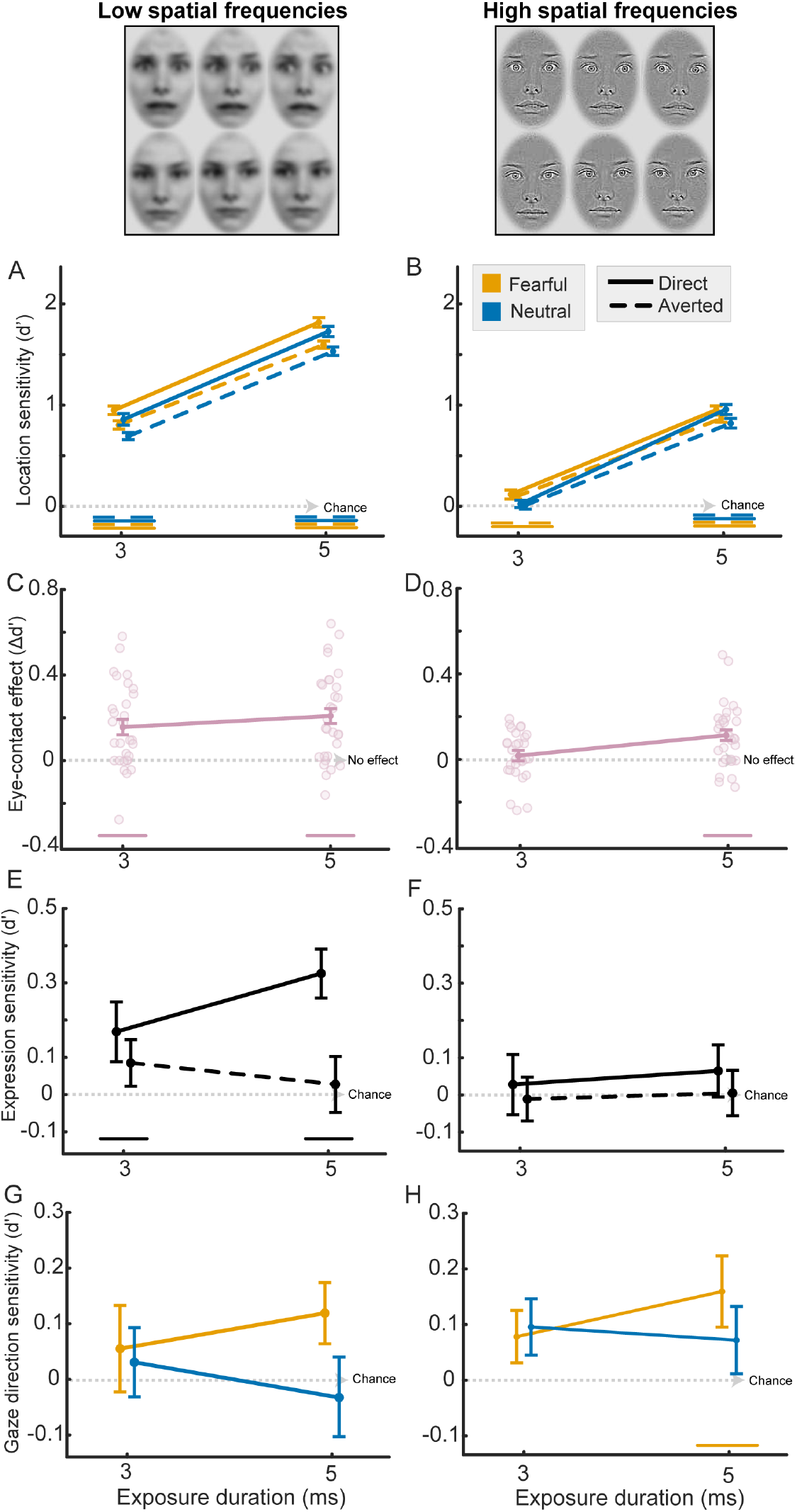
Type-1 sensitivity across spatial frequency conditions in Experiment 2. **(A**,**B)** Localisation sensitivity for **(A)**, low-spatial-frequency, LSF, and **(B)**, high-spatial-frequency, HSF, faces, indexed by type-1 *d*′ for detecting the location of the intact face. Sensitivity increased with exposure duration, was higher for LSF than HSF faces, and was higher for direct-gaze than averted-gaze faces. **(C**,**D)** Eye-contact effect, indexed as the direct-minus-averted difference in localisation sensitivity, for **(C)**, LSF, and **(D)**, HSF, faces. The direct-gaze advantage was evident in the LSF condition at both exposures, but emerged in the HSF condition only at 5 ms. **(E**,**F)** Expression sensitivity for **(E)**, LSF, and **(F)**, HSF, faces, indexed by type-1 *d*′ for categorising fearful versus neutral facial expressions. Expression sensitivity was reliable only for direct-gaze faces in the LSF condition. **(G**,**H)** Gaze direction sensitivity for **(G)**, LSF, and **(H)**, HSF, faces, indexed by type-1 *d*′ for categorising direct versus averted gaze. Gaze direction sensitivity was comparatively weak, with only limited evidence of above-chance performance. Horizontal lines above the x-axis indicate exposure durations with above-chance sensitivity, or direct-minus-averted differences above zero, after Holm correction. Data are presented as mean *±*1 SEM. All face stimuli, including the intact face shown here, were taken from the Radboud Face Database (RaFD) and are presented as a stimulus example (see: https://rafd.nl/)

To characterise the gaze-direction effect more directly, we examined the eye-contact effect as the direct-minus-averted difference in localisation *d*′ (Fig. 6C,D). In the LSF condition, the direct-gaze advantage exceeded zero for both fearful and neutral faces at both exposure durations (all Holm-corrected *p ≤* 0.005). In the HSF condition, by contrast, the direct-gaze advantage was absent at 3 ms for both expressions (both Holm-corrected *p ≤* 0.309), but emerged at 5 ms for both fearful and neutral faces (both Holm-corrected *p ≤*0.043). Thus, the eye-contact effect was already reliable under coarse spatial filtering at 3 ms, whereas under fine spatial filtering it became reliable only at the longer exposure.

Expression sensitivity showed a more restricted pattern (Fig. 6E,F). Although expression sensitivity was higher for direct-gaze than averted-gaze faces overall (*F* _(1,29)_ = 4.27, *p* = 0.048, 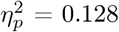), condition-wise tests showed that expression sensitivity was reliably above zero only for direct-gaze LSF faces, at both 3 and 5 ms. All other conditions were non-significant after correction. Thus, explicit expression categorisation was supported only when direct gaze was available in coarse spatial information.

Gaze-direction sensitivity was weak overall (Fig. 6G,H). The omnibus analysis showed no significant main effects of expression, spatial frequency or exposure duration, and condition-wise tests showed only limited evidence of above-chance gaze categorisation after correction. Thus, unlike localisation, explicit gaze-direction categorisation remained unreliable or weak under ultra-brief spatial-frequency-filtered presentation.

Together, these findings indicate that coarse spatial information supported robust localisation and the earliest eye-contact effect, whereas HSF content supported a direct-gaze advantage only at the longer exposure. Explicit expression and gaze categorisation remained limited. See Supplementary Notes 5 and 6 for details.

### Information-theoretic analyses confirm that coarse spatial information supports localisation

We next asked whether the spatial-frequency pattern observed in signal detection sensitivity was also present when performance was expressed as stimulus-response information. Localisation responses carried reliable information about stimulus location for LSF faces at both 3 and 5 ms, across all four expression-by-gaze conditions (Fig. 7A). For HSF faces, by contrast, localisation information was not reliably above zero at 3 ms, but became reliable in all conditions at 5 ms (Fig. 7B). Thus, localisation information emerged earlier and more strongly for LSF than HSF faces.

**Figure 7:**
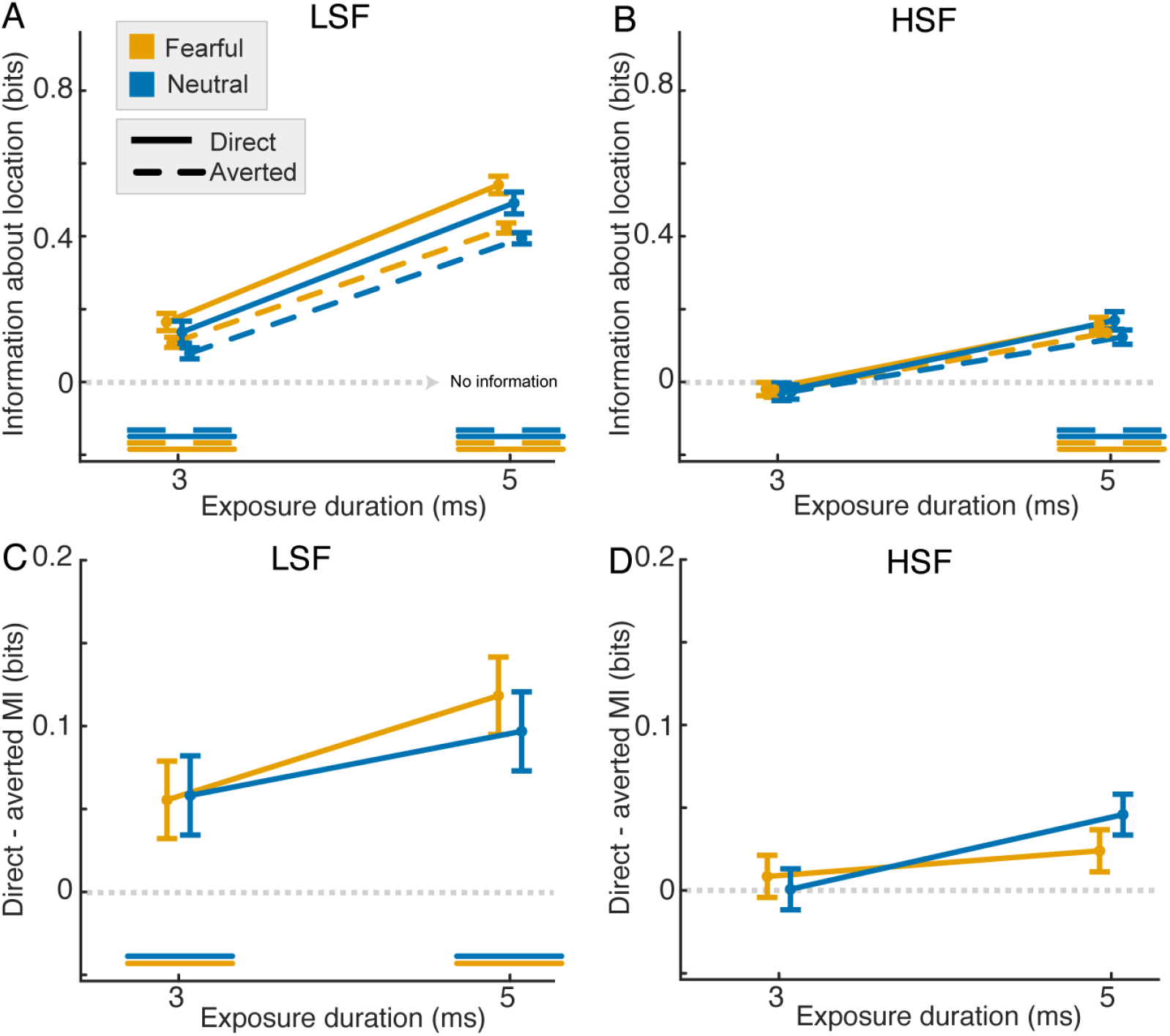
Information-theoretic evidence that low-spatial-frequency information supports the eye-contact effect in Experiment 2. **(A**,**B)** Mutual information between the actual location of the intact face and participants’ localisation responses, *I*(stimulus location; localisation response), shown separately for **(A)**, low-spatial-frequency, LSF, and **(B)**, high-spatial-frequency, HSF, faces. Localisation responses carried reliable information about stimulus location for LSF faces at both 3 ms and 5 ms, whereas HSF localisation information became reliable only at 5 ms. **(C**,**D)** Direct-gaze advantage in localisation information, computed as the direct-minus-averted difference in localisation mutual information, shown separately for **(C)**, LSF, and **(D)**, HSF, faces. Direct gaze increased LSF localisation information at 3 ms for fearful faces and at 5 ms for both fearful and neutral faces. HSF direct-minus-averted contrasts did not survive correction. Lines indicate group means and error bars indicate within-subject SEM. Dotted horizontal lines indicate zero information or zero contrast. Horizontal markers indicate exposure durations at which mutual information was significantly greater than zero after Holm correction.

Direct gaze increased localisation information most clearly in the LSF condition (Fig. 7C,D). For LSF faces, direct-gaze stimuli carried more localisation information than averted-gaze stimuli at 3 ms for fearful expressions, and at 5 ms for both fearful and neutral expressions. In the HSF condition, direct-minus-averted localisation information contrasts did not survive correction at either exposure duration. These information-theoretic results converged with the signal-detection analyses in showing that coarse visual information provided the clearest support for early eye-contact-related localisation. See Supplementary Note 7 for additional results.

### Autistic traits were associated with a reduced eye-contact effect on localisation sensitivity

Finally, we examined whether autistic traits, measured with the Autism-spectrum Quotient (AQ), predicted individual differences in gaze- and expression-related sensitivity effects in Experiment 1. Higher AQ scores were associated with a smaller eye-contact effect on localisation sensitivity when direct-minus-averted contrast scores were averaged across exposure durations (two-sided Pearson correlation, r = −0.396, 95% CI [−0.662, −0.042], t(28) = −2.285, *p* = 0.03; Fig. 8A). This association was not observed for the eye-contact effect on metacognitive sensitivity, the fearful-neutral difference in localisation sensitivity, or the fearful-neutral difference in metacognitive sensitivity (Fig. 8B,D). Thus, autistic traits were associated with a reduced objective localisation advantage for direct gaze over averted gaze, but not with metacognitive sensitivity to eye contact or with localisation or metacognitive effects of emotional expression. This association should be interpreted cautiously, as it did not survive correction when the four Figure 8 correlations were treated as a single family of tests.

**Figure 8:**
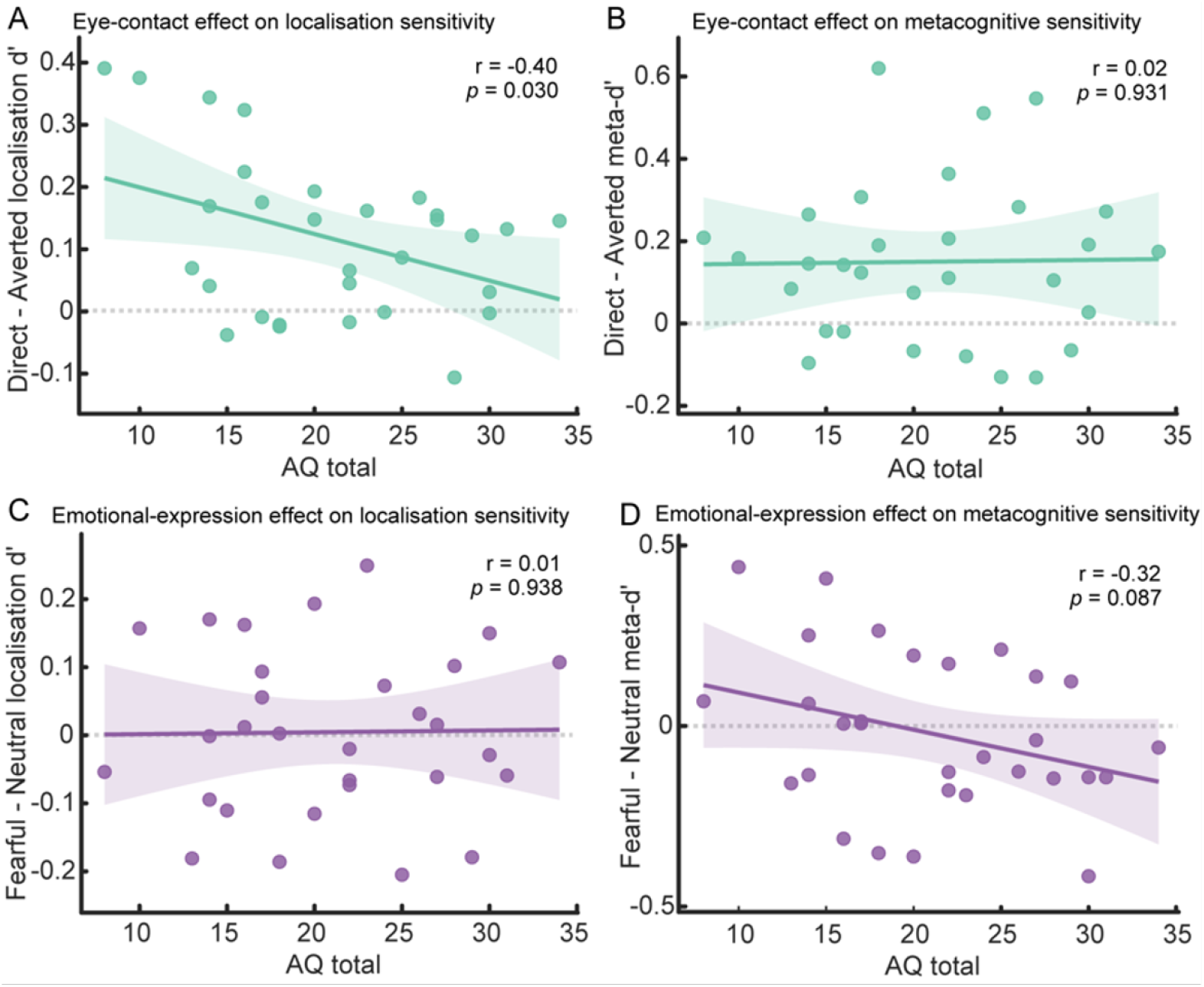
Autistic traits and gaze- or expression-related sensitivity effects in Experiment 1. **(A)** Eye-contact effect on localisation sensitivity, indexed as the difference in type-1 *d*′ between direct-gaze and averted-gaze faces. Higher Autism-spectrum Quotient, AQ, scores were associated with a smaller localisation advantage for direct gaze over averted gaze, Pearson’s *r* = *−* 0.396, 95% CI [*−* 0.662, *−* 0.042], *t*(28) = *−* 2.285, *p* = 0.030; Holm-corrected *p* across the four Figure 8 correlations = 0.120. Eye-contact effect on metacognitive sensitivity, indexed as the difference in meta-*d*′ between direct-gaze and averted-gaze faces. AQ did not predict this metacognitive eye-contact effect, *r* = 0.017, 95% CI [*−* 0.346, 0.375], *t*(28) = 0.088, *p* = 0.931. **(C)** Fearful-neutral difference on localisation sensitivity, indexed as the difference in type-1 *d*′ between fearful and neutral faces. AQ did not predict this fearful-neutral expression difference on localisation sensitivity, *r* = 0.015, 95% CI [ *−*0.347, 0.373], *t*(28) = 0.079, *p* = 0.938. **(D)** Fearful-neutral difference on metacognitive sensitivity, indexed as the difference in meta-*d*′ between fearful and neutral faces. AQ did not reliably predict this fearful-neutral difference on metacognitive sensitivity, *r* = 0.317, 95% CI [ *−*0.608, 0.048], *t*(28) = *−*1.771, *p* = 0.087. For all panels, contrast scores were averaged across exposure durations before correlation with AQ. Positive values indicate greater sensitivity for direct than averted gaze in Panels A and B, and greater sensitivity for fearful than neutral expressions in Panels C and D. Dots represent individual participants; lines show least-squares regression fits with shaded 95% confidence intervals. Pearson correlations were two-sided.

## DISCUSSION

Eye contact is among the most salient signals in human social perception, but it has remained unclear how much visual information is required for the visual system to register that another individual is looking at us, and whether this registration requires conscious awareness. Across two experiments, direct gaze facilitated objective face localisation at ultra-brief exposure durations, before observers could reliably categorise gaze direction and before subjective experience reliably tracked the perceptual evidence supporting performance. Information-theoretic analyses complemented this pattern by showing that direct gaze increased stimulus-location information in objective localisation responses, whereas explicit categorisation and subjective awareness became informative later. This early eye-contact effect depended primarily on LSF information, whereas HSF information contributed more clearly at longer exposures. Finally, autistic traits in neurotypical adults were associated with a reduced eye-contact effect on localisation sensitivity, although this association should be interpreted cautiously. Together, these findings suggest that the visual system can register socially relevant gaze information from remarkably sparse sensory input, before this information becomes available to conscious experience.

Direct gaze reduced the visual input required for face localisation. Psychometric estimates showed lower criterion thresholds for direct than averted gaze, and signal detection analyses showed a localisation advantage from 3 ms onwards, before gaze categorisation was reliable. Information theory analyses showed the same effect as increased stimulus location information in objective responses. Thus, gaze direction influenced face localisation before participants could accurately report whether the eyes were direct or averted. These findings extend previous evidence that direct gaze facilitates access to awareness under interocular suppression[13, 14, 42–44] by showing that eye contact prioritisation can be observed under precisely controlled ultra-brief stimulation, without masking or suppression paradigms.

The facial-expression manipulation provided an important specificity test. Fearful expressions have often been reported to gain preferential access to awareness under bCFS[45], but such findings are difficult to interpret because breakthrough times can reflect low-level image differences, response criteria or other post-perceptual processes rather than perceptual sensitivity itself[17, 46]. Subsequent work has likewise shown that apparent fear advantages can depend on low-level visual properties and spatial-frequency content, rather than emotional expression per se[46–49]. Consistent with this more cautious view, emotional expression did not reliably modulate localisation thresholds, localisation sensitivity or metacognitive sensitivity in the present study. The information-theoretic analyses reinforced this specificity: expression and gaze categorisation responses carried comparatively little information at the shortest exposures, and the strongest stimulus-response information was observed for localisation rather than explicit categorisation. Thus, the early effect observed here appears to reflect a selective advantage for direct gaze rather than a broader prioritisation of socially salient facial features.

A central challenge for interpreting detection advantages is that they may reflect either unconscious processing of the relevant stimulus dimension or partial conscious access to that dimension. To address this challenge, we adapted the detection-discrimination dissociation logic[37] by adding a metacognitive-access criterion. Under this framework, evidence for unconscious processing is strongest when a stimulus dimension influences detection, while explicit discrimination of that dimension remains at chance and subjective ratings do not reliably distinguish correct from incorrect responses. The present results approximate this pattern at the shortest exposure durations. Direct gaze enhanced localisation sensitivity before gaze categorisation became reliable, and before metacognitive sensitivity indicated reliable access to the perceptual evidence supporting localisation. This suggests that the early direct-gaze advantage was not accompanied by reliable conscious access, either in the form of explicit gaze-direction categorisation or in the form of metacognitive insight into performance.

The metacognitive and information-theoretic results clarified this dissociation. Localisation sensitivity was reliable from 2 ms onwards, and the direct-gaze advantage emerged from 3 ms onwards. By contrast, metacognitive sensitivity and metacognitive efficiency became reliable only at longer exposure durations, from approximately 4 ms onwards. Consistent with this pattern, co-information between stimulus location, localisation response and PAS ratings became reliably positive only at the longest exposure duration for most expression-by-gaze conditions. Thus, participants could use visual evidence to localise the intact face before their subjective experience reliably tracked, overlapped with, or modulated that localisation information. This suggests that eye contact first biases objective localisation outside reliable awareness, and later enhances the mapping between perceptual evidence and subjective experience.

Experiment 2 provided converging evidence that the earliest eye-contact effect was supported primarily by coarse visual information. LSF faces produced robust localisation performance and a reliable direct-gaze advantage at both 3 and 5 ms. Information-theoretic analyses confirmed this pattern by showing that localisation responses carried reliable information about stimulus location for LSF faces at both exposure durations, whereas HSF localisation information became reliable only at 5 ms. Direct gaze also increased localisation information most clearly in the LSF condition. Thus, the earliest stages of eye-contact processing appear to rely primarily on coarse visual structure, whereas fine-grained visual information contributes more clearly once additional sensory evidence is available. This pattern is consistent with accounts proposing that eye contact can be initially detected through rapid processing routes before modulating cortical social brain networks[1, 33–36]. However, spatial-frequency manipulations alone cannot establish subcortical involvement. Future work using temporally precise neural measures, causal perturbation or neuroimaging will be needed to determine whether early eye-contact effects depend on subcortical pathways, early visual cortex, face-selective cortical regions or interactions among these systems.

The spatial-frequency findings also suggest that direct gaze may influence other aspects of face processing at minimal exposure durations. In Experiment 2, expression sensitivity was reliable only for direct-gaze faces in the LSF condition, whereas explicit gaze-direction sensitivity remained weak overall. This pattern is consistent with the idea that direct gaze can enhance the processing of socially relevant facial information even when explicit categorisation of gaze direction is limited. However, the expression effect was more restricted than the localisation advantage and should therefore be interpreted cautiously. Thus, the primary conclusion is not that all aspects of social face perception are available under minimal visual input, but rather that direct gaze selectively facilitates early objective localisation through coarse visual information.

Autistic traits were associated with individual differences in the early eye-contact effect. Participants with higher AQ scores showed a smaller direct-gaze advantage in localisation sensitivity when contrast scores were averaged across exposure durations. This association was selective: AQ did not predict the eye-contact effect on metacognitive sensitivity, nor did it predict fearful-neutral differences in localisation or metacognitive sensitivity. This pattern is consistent with the possibility that autistic traits are linked to reduced perceptual prioritisation of eye contact at early stages of visual processing[50–54], rather than to a broad reduction in sensitivity to facial social cues or a general alteration in metacognitive access. This finding should be interpreted cautiously because the sample was neurotypical, the association did not survive correction across the four correlations, and the study was not designed as a clinical comparison.

More broadly, these findings may help clarify the function of conscious awareness in social perception. In our data, direct gaze influenced objective localisation before observers could reliably categorise gaze direction and before subjective awareness reliably tracked the evidence supporting performance. Information-theoretic analyses sharpened this conclusion by showing that objective responses carried stimulus-location information before PAS ratings became reliably coupled to that information. This suggests that conscious awareness may not be necessary for the earliest prioritisation of socially relevant signals, but may become important when those signals must be stabilised into explicit, reportable and metacognitively accessible representations. The present results therefore fit with proposals that some stimulus dimensions can bias detection outside awareness, while conscious access is required for richer perceptual discrimination and flexible use of information[55]. They also suggest an interesting contrast with bodily self-perception[56–58], where recent work indicates a much tighter coupling between perceptual processing and conscious access[59].

In summary, direct gaze facilitated face localisation at the limits of visual stimulation, before explicit gaze categorisation and before reliable metacognitive access to the relevant perceptual evidence. Information-theoretic analyses confirmed that this effect reflected increased stimulus-location information in objective localisation responses, and that subjective awareness became coupled to this information only later. This early eye-contact effect was supported primarily by coarse spatial information and was reduced in individuals with higher autistic traits. These findings suggest that eye contact is prioritised at an exceptionally early stage of visual processing, using minimal sensory input that can influence behaviour even when exposure durations are insufficient for conscious access to gaze direction.

## METHODS

### Participants

The experiments were approved by the Ethics Committee of the Faculty of Psychological Science and Education at the Université libre de Bruxelles. All participants provided informed consent and received €15 per hour for participating in Experiments 1 and 2.

Thirty-three participants were recruited to take part in both experiments. Three participants were excluded from the analyses: one because they failed to respond on more than 5% of trials, one because their accuracy was at chance across all exposure durations, suggesting that they were not engaged with the task, and one because they asked to stop the experiment halfway through. This left a final sample of 30 participants (M_age_ = 25.72 years, SD_age_ = 11.14; 17 female).

Participants self-reported their age and gender. These variables were not considered in the study design or included in the analyses, as no hypotheses concerned age or gender. All participants had normal or corrected-to-normal vision and reported no history of neurological or psychiatric disorders.

### Display and apparatus

The custom-made LCD tachistoscope used in this study was based on the design described by Sperdin et al.[23, 60]. The apparatus consisted of two LCD screens: one positioned vertically and the other horizontally, aligned with the top of the vertical screen. A semi-permeable mirror was placed diagonally between the two screens, allowing light from the vertical screen to pass through while reflecting light from the horizontal screen. As a result, when the backlights of both screens were switched on simultaneously, the observer saw the two displays superimposed. The setup allowed precise control over which screen was visible to the observer by regulating the screens’ backlights. These were controlled by a dedicated microcontroller with a temporal precision of 0.002 *±*0.001 ms. In all experiments, the vertical screen presented all images that were not displayed for ultra-brief durations below 10 ms, including fixation crosses, placeholders, and response cues; consequently, its backlight remained continuously on.

The horizontal screen presented the stimuli shown for durations below 10 ms, and its backlight remained off except during these brief stimulus presentations. During these ultra-brief intervals, the content of both screens was visible to the observer. Full technical details of the apparatus are provided in Supplementary Note 1 in Lanfranco et al.[15]

### Stimuli

Face stimuli were selected from the Radboud Faces Database[61] (RaFD). We selected six identities (3 female, 3 male) that were closely matched on the database’s normative ratings of expression recognisability, emotional intensity and arousal. For each identity, we used fearful and neutral expressions. Within each expression category, half of the stimuli displayed direct gaze and half displayed averted gaze, with averted-gaze stimuli distributed across leftward and rightward deviations (see Supplementary Fig. 1).

All images were pre-processed using a common pipeline (Extended Data Fig. 1). First, the original photographs were cropped to retain the internal facial region and converted to greyscale. Scrambled control images were then created by Fourier phase scrambling, such that the phase spectrum was randomised while the amplitude spectrum was preserved. Intact and scrambled images were subsequently equated for mean luminance using the SHINE toolbox[62, 63]. After luminance matching, each image was presented within an oval aperture on a uniform grey background. The transition between the face region and the background was softened by applying a Gaussian-blurred mask, reducing sharp edge cues. Final stimuli were resized to a common image size before presentation.

For Experiment 2, we additionally generated spatial-frequency-filtered versions of the intact and scrambled stimuli. LSF and HSF images were created by applying Gaussian low-pass and high-pass filters in the Fourier domain, respectively. The filter settings corresponded approximately to coarse and fine spatial scales, allowing us to compare the contribution of lower and higher spatial-frequency information to performance. After filtering, these images were again placed within the blurred oval aperture and matched to the same output format as the broadband stimuli.

### Analysis

#### Psychometric function analysis

To quantify the minimum visual input required for face localisation, we fitted psychometric functions to performance in the two-alternative forced-choice location task. For each participant, we computed the proportion of correct localisation responses at each exposure duration (1– 5 ms), separately for the four face conditions defined by emotional expression and gaze direction (fearful-direct, fearful-averted, neutral-direct and neutral-averted). These condition-wise accuracy values were then fitted with a sigmoidal psychometric function relating exposure duration, *x*, to the probability of a correct response, Ψ(*x*). At the group level, we visualised both the fitted mean psychometric functions and the corresponding individual participant fits.

In general form, the fitted psychometric function can be written as

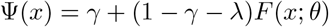

where *γ* is the guess rate, *λ* is the lapse rate, and *F* (*x*; *θ*) is a monotonic sigmoid with parameters *θ* governing its location and steepness. Because the localisation task was a two-alternative forced-choice task, the lower asymptote was constrained by chance performance, such that *γ* = 0.5. The fitted sigmoid was implemented as a cumulative Gaussian function, written as

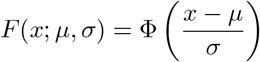

where Φ denotes the standard normal cumulative distribution function, *µ* indexes the central tendency of the function, and *σ* its spread. Under this parameterisation, the exposure duration corresponding to 75% correct performance was obtained by solving

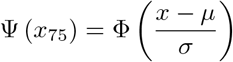

which yields

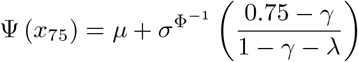

This value was taken as the threshold at 75% correct (*thr75*), with lower values indicating that less visual input was required to achieve criterion performance.

From each fitted function, we extracted three summary measures. First, we estimated the threshold at 75% correct (*thr75*), which indexed the minimum exposure duration required to reach criterion-level localisation performance. Second, we derived the just-noticeable difference (JND) from the fitted psychometric function as an index of the sharpness of the transition from chance to asymptotic performance, with smaller values indicating greater precision. For symmetric psychometric functions, JND can be expressed as

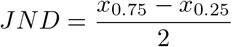

where *x*_*p*_ denotes the stimulus value associated with response probability *p* after accounting for the fitted asymptotes. Smaller JND values indicate a steeper and more precise transition from chance to asymptotic performance. Third, we extracted the slope parameter, *β*, from the fitted psychometric function as a complementary index of steepness, such that larger *β* values correspond to more abrupt changes in performance with increasing exposure duration. These three measures were then carried forward to the group-level analyses.

### Type-1 signal-detection analysis

To dissociate bias-independent sensitivity from response bias, we quantified type-1 signal detection measures[24, 25] separately for the location, expression and gaze tasks. All measures were computed separately for each participant and condition, and then entered into the group-level analyses.

For the location task, participants indicated whether the intact face had appeared on the left or right side of the display. Because this was a two-alternative forced-choice judgement, location sensitivity was estimated as

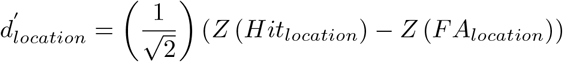

where *Hit* denotes the hit rate, defined as the proportion of trials on which a face presented on the right was correctly reported as appearing on the right, and *FA* denotes the false-alarm rate, defined as the proportion of trials on which a face presented on the left was incorrectly reported as appearing on the right. Response criterion for the location task was computed as

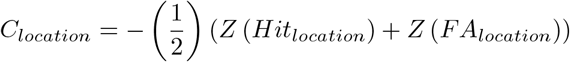

Positive values indicate a tendency to respond “left”, whereas negative values indicate a tendency to respond “right”.

For the expression task, participants categorised each face as fearful or neutral. Expression sensitivity was estimated using the standard yes/no SDT formulation,

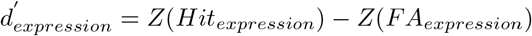

where hits were defined as trials on which a fearful face was correctly classified as fearful, and false alarms were defined as trials on which a neutral face was incorrectly classified as fearful. Response criterion for expression judgements was calculated as

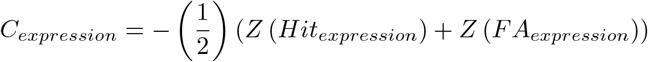

For the gaze task, participants categorised each face as showing direct or averted gaze. Gaze-direction sensitivity was likewise estimated using the standard yes/no SDT formulation,

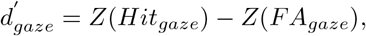

where hits were defined as trials on which direct gaze was correctly identified as direct, and false alarms were defined as trials on which averted gaze was incorrectly classified as direct. Response criterion for gaze-direction judgements was computed as

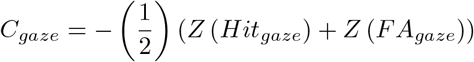

Where necessary, extreme hit and false-alarm rates were adjusted using a standard edge-correction procedure before *Z*-transformation to avoid infinite values. In Experiment 1, these measures were computed separately for each display duration and stimulus condition. In Experiment 2, the same procedure was used, with spatial frequency included as an additional within-subject factor.

### Type-2 signal-detection analysis (maximum-likelihood estimation)

To quantify how closely subjective awareness tracked objective performance, we estimated type-2 SDT measures from participants’ Perceptual Awareness Scale (PAS) ratings using the maximum-likelihood procedure developed by Maniscalco and Lau[28]. These analyses were based on the location task, such that metacognitive sensitivity indexed the extent to which PAS ratings discriminated between correct and incorrect localisation responses.

For each participant and condition, trials were sorted according to type-1 stimulus class, type-1 response accuracy, and PAS rating. These counts were then arranged into the response-frequency vectors required for type-2 SDT, *nR*_*S*1_ and *nR*_*S*2_, which summarise the distribution of awareness ratings conditional on the two type-1 stimulus classes. We then fitted the equal-variance SDT model by maximum likelihood to estimate meta-*d*′, which reflects the amount of type-1 signal that would be required for an ideal observer to produce the observed pattern of type-2 responses. Higher meta-*d*′ values therefore indicate that subjective awareness ratings more closely tracked objective discrimination performance.

From the same model, we also derived metacognitive efficiency and metacognitive bias. Metacognitive efficiency was quantified as

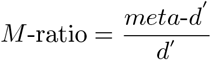

where *d*′ was the corresponding type-1 sensitivity estimate from the location task. Values close to 1 indicate that metacognitive sensitivity closely matched the amount of information available for type-1 performance, whereas values below 1 indicate reduced metacognitive efficiency.

Metacognitive bias was indexed by the placement of type-2 criteria, reflecting the tendency to use higher or lower awareness ratings independently of objective performance. All type-2 SDT measures were estimated separately for each participant and condition and were subsequently analysed at the group level.

### Hierarchical metacognitive efficiency analysis

Because individual-participant estimates of metacognitive efficiency can be unstable when trial counts are not abundant, we also estimated metacognitive efficiency using the hierarchical Bayesian HMeta-*d*′ model[29]. This approach jointly estimates participant-level and group-level parameters under partial pooling, thereby improving the stability of metacognitive estimates while preserving between-participant variability.

For each condition, participant-specific *nR*_*S*1_ and *nR*_*S*2_ response-count vectors derived from PAS ratings in the location task were entered into the hierarchical model. The model estimates posterior distributions over meta-*d*′ and *M*-ratio at both the individual and group levels. As in the maximum-likelihood analysis, *M*-ratio was defined as meta-*d*′/*d*′, with higher values indicating more efficient metacognitive access relative to type-1 sensitivity. Group-level posterior distributions were summarised using their central tendency and 95% highest density intervals (HDIs). To compare conditions, we computed posterior contrasts by subtracting posterior samples from the relevant group-level parameter estimates. A condition difference was interpreted as credible when the 95% HDI of the posterior contrast excluded zero. This hierarchical analysis was used primarily to quantify condition differences in metacognitive efficiency across exposure durations, while taking into account the limited number of trials contributing to participant-level type-2 estimates.

### Information-theoretic analysis

To complement the signal-detection analyses, we quantified the information carried by behavioural responses about the task-relevant stimulus variables using discrete information-theoretic measures[31, 64]. Mutual information was estimated in bits from empirical joint probability distributions computed at the single-participant level. For two discrete variables *X* and *Y*, mutual information was defined as

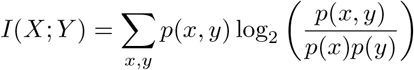

For the localisation task, *X* was the actual location of the intact face and *Y* was the participant’s localisation response. Thus, *I* (stimulus location; localisation response) quantified how much information participants’ left-right responses carried about the true location of the intact face. These estimates were computed separately for each participant, exposure duration, emotional expression and gaze direction. For the expression task, *X* was the emotional expression shown and *Y* was the expression response; estimates were computed separately for each participant, exposure duration and gaze direction. For the gaze-direction task, *X* was the gaze direction shown and *Y* was the gaze response; estimates were computed separately for each participant, exposure duration and emotional expression. To assess whether direct gaze increased localisation information, we computed participant-level planned contrasts by subtracting localisation mutual information for averted-gaze faces from localisation mutual information for direct-gaze faces, separately for each exposure duration and emotional expression. Analogous specificity contrasts were computed for expression information, as the direct-minus-averted difference in *I* (expression; expression reported), and for gaze information, as the fearful-minus-neutral difference in *I* (gaze direction; gaze direction reported).

To relate localisation information to subjective awareness, we computed three-variable co-information between stimulus location, localisation response, and PAS ratings. Co-information was defined as

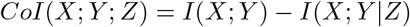

where *X* was the actual stimulus location, *Y* was the localisation response, and *Z* was the PAS rating. Here, *CoI* (stimulus location; localisation response; PAS) quantified whether subjective visibility ratings overlapped with, or modulated, the information shared between the actual stimulus location and the localisation response. These estimates were computed separately for each participant, exposure duration, emotional expression and gaze direction. Positive values indicate that PAS ratings accounted for shared stimulus-response information, whereas negative values indicate that conditioning on PAS revealed additional stimulus-response dependency. Because co-information is a signed interaction measure[65], it was not interpreted as a direct estimate of redundancy or synergy. All information-theoretic estimates were computed from discrete trial-level variables, with invalid or missing trials excluded. PAS ratings were treated as a four-level discrete variable. No binning was applied to the behavioural response variables because stimulus, response and PAS variables were already discrete. To reduce finite-sample bias in plug-in information estimates, each observed value was bias-corrected by subtracting the mean of a permutation-based null distribution[32, 66]. For mutual information, the null distribution was obtained by randomly permuting the relevant stimulus labels within each participant and analysis cell. For PAS co-information, PAS labels were permuted while preserving the pairing between stimulus location and localisation response. We used 1,000 permutations per estimate. Bias-corrected values could therefore fall slightly below zero; such values were interpreted as no evidence for information above the permutation baseline rather than as negative information.

Group-level inference was performed on participant-level bias-corrected estimates. Condition-wise mutual information and PAS co-information values were tested against zero using one-sample t-tests. Planned contrast values were also tested against zero using one-sample t-tests. *p* values were corrected for multiple comparisons using the Holm procedure within each family of tests. Effect sizes are reported as Cohen’s d_z_. Data are plotted as group means with within-subject SEM.

### Correlation analyses

To test whether autistic traits are associated with a delayed eye-contact effect in the 1-5 ms-exposure range, we conducted correlation analyses relating autistic traits to each contrast measure. AQ total scores were correlated with four participant-level contrast variables: the direct-minus-averted difference in location *d*′, the direct-minus-averted difference in meta-*d*′, the fearful-minus-neutral difference in location *d*′, and the fearful-minus-neutral difference in meta-*d*′. These contrasts were computed separately for each participant at each exposure duration. Direct-minus-averted contrasts were calculated after collapsing across emotional expression, whereas fearful-minus-neutral contrasts were calculated after collapsing across gaze direction.

For each contrast measure, we first computed exposure-specific correlations between AQ total and contrast strength across participants. We report both Pearson product-moment correlations and Spearman rank correlations, to assess linear and monotonic associations, respectively. These analyses were conducted separately at each exposure duration. To control for multiple comparisons across exposure-specific tests, *p* values were Holm-corrected within each contrast measure, separately for Pearson and Spearman correlations.

We also computed an overall correlation for each contrast measure by first averaging contrast values across exposure durations within each participant and then correlating this participant-level mean contrast with AQ total. These overall associations were likewise assessed using both Pearson and Spearman coefficients. These correlation analyses were treated as complementary to the mixed-effects models and were used to characterise the association between autistic traits and the magnitude of gaze-related and emotion-related contrast effects.

### Statistical information

Statistical analyses were conducted using repeated-measures and mixed-effects approaches, depending on the structure of the data. Unless otherwise specified, the significance threshold was set at *α* = 0.05. For the main behavioural and metacognitive analyses in Experiments 1 and 2, we used repeated-measures analyses of variance (ANOVAs) with participant as the repeated factor. These models were fitted to complete-case datasets, such that only participants with data in all cells relevant to a given analysis contributed to that model. For omnibus repeated-measures ANOVAs, Greenhouse–Geisser-corrected *p* values were reported when available. Effect sizes for ANOVAs are reported as partial eta squared 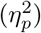.

When significant interactions were observed, these were followed by planned paired-samples *t*-tests within the relevant factor levels. We also used one-sample *t*-tests against zero to assess whether sensitivity or contrast measures exceeded chance-level performance. These tests were one-tailed for sensitivity measures and two-tailed for bias measures. Effect sizes for paired and one-sample *t*-tests are reported as Cohen’s *d* z. Across follow-up and against-zero analyses, *p* values were corrected for multiple comparisons using the Holm procedure within each relevant family of tests.

For Experiment 1, repeated-measures analyses were applied to type-1 sensitivity and bias measures for localisation, expression categorisation and gaze-direction categorisation, as well as to type-2 measures of metacognitive sensitivity and metacognitive efficiency. For Experiment 2, analogous repeated-measures ANOVAs were used, with spatial frequency added as a within-subject factor. Psychometric-function-derived summary measures, including the threshold at 75% correct (*thr75*), the just-noticeable difference (JND), and the slope parameter (*β*), were analysed at the group level using repeated-measures designs matched to the factorial structure of each experiment.

To examine associations between autistic traits and contrast effects in Experiment 1, we used linear mixed-effects models of the form Contrast *~z*AQ*×* Exposure + (1 | Participant), where AQ total scores were standardised before analysis. Separate models were fitted for the direct-minus-averted differences in location *d*′ and meta-*d*′, and for the fearful-minus-neutral differences in location *d*′ and meta-*d*′. These models tested whether AQ predicted contrast strength overall and whether that association varied across exposure durations, while accounting for repeated observations within participant through a random intercept. Denominator degrees of freedom and *p* values were estimated using the Satterthwaite approximation. To complement these models, we also computed exposure-specific Pearson and Spearman correlations between AQ total and each contrast measure, as well as participant-level correlations using contrast values averaged across exposure durations. Holm correction was applied across exposure-specific correlations within each contrast family, separately for Pearson and Spearman analyses.

Data are reported as mean *±*SEM unless otherwise stated. All tests were two-sided unless a directional hypothesis justified one-tailed against-zero tests for sensitivity measures.

## DATA AVAILABILITY

The data supporting the findings of this study will be made publicly available via the Open Science Framework (OSF) after publication.

## CODE AVAILABILITY

The custom code supporting the analyses reported in this study will be made publicly available via the OSF after publication.

## ACKNOWLEDGEMENTS

R.C.L. was supported by the Swedish Research Council (grant number 2024-00839), the Strategic Research Area Neuroscience (StratNeuro), the National Research and Development Agency-Chile (grant number #74220010), and a ULB Seal of Excellence Research fellowship. A.G. was supported by the European Research Council (grant number 101220685-MINDSIM), Swedish Research Council (grant number 2021-02089), the Swedish Foundation for Strategic Research (grant number FFL21-0261), the Wenner-Gren Foundations (grant number FT2021-0003), the Swedish Society of Medicine (grant number SLS-960489), the Swedish Brain Foundation (grant number FO2024-0385), the Swedish Society for Medical Research (grant number SG-24-0184-B-H-01), and the Marcus and Amalia Wallenberg Foundation (grant number 2024.0042). A.C. was supported by the European Research Council (grant number 101055060-EXPERIENCE).

## AUTHOR CONTRIBUTIONS

R.C.L. and A.C. conceived the study. R.C.L. conceptualised the study. R.C.L. and A.C. acquired funding. R.C.L. designed the experiments, programmed the experiments, collected the data, created the code, analysed the data, and created the figures. R.C.L. wrote the manuscript with input from A.G. and A.C. Finally, R.C.L., A.G., and A.C. edited the manuscript.

