## Supplementary Note for "Conscious and unconscious eye contact at the limits of vision"

### EXPERIMENT 1

#### Supplementary Note 1. Stimulus pre-processing and spatial-frequency filtering

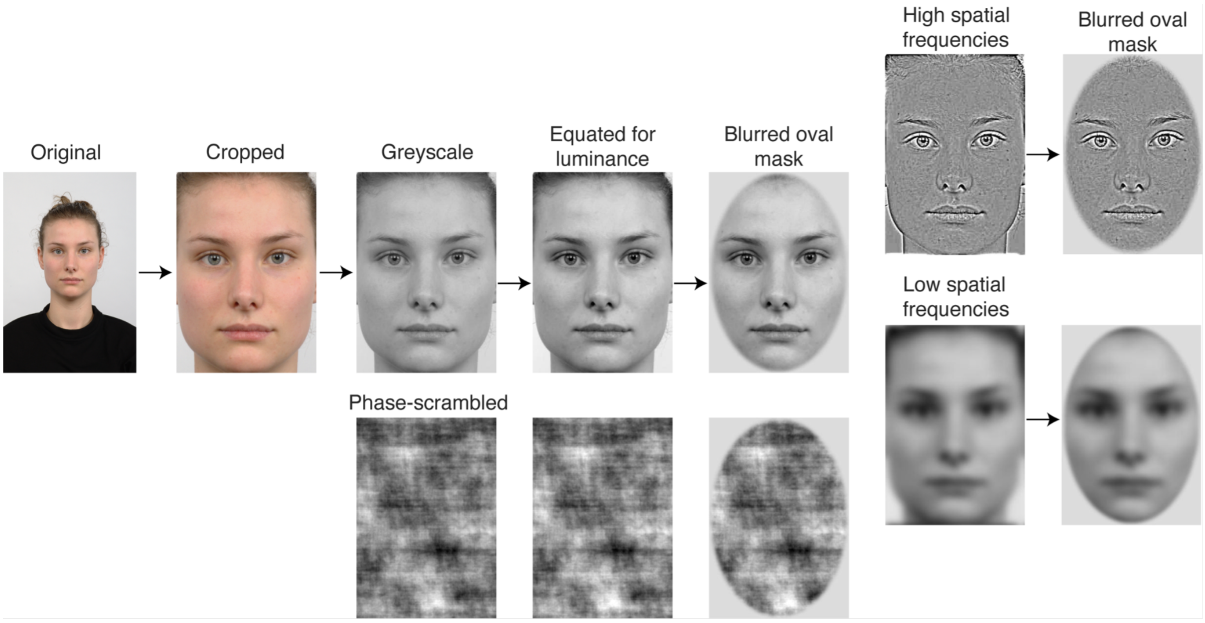


**Supplementary Fig. 1. Stimulus pre-processing and spatial-frequency filtering.** Representative example of the stimulus pre-processing pipeline. Original face images were cropped to retain the internal facial region, converted to greyscale, equated for luminance, and presented within a blurred oval aperture to reduce sharp edge cues. Phase-scrambled control images were generated by randomising the phase spectrum while preserving the amplitude spectrum of the corresponding face image, and were processed using the same luminance-equating and masking steps. For Experiment 2, the final masked face images were additionally filtered to create high-spatial-frequency and low-spatial-frequency versions, allowing us to test whether fine-grained or coarse visual information preferentially supported eye-contact processing at ultra-brief exposure durations. The same preprocessing pipeline was applied to all identities, expressions and gaze-direction conditions. All face stimuli, including the intact face shown here, were taken from the Radboud Face Database (RaFD) and are presented as a stimulus example (see: https://rafd.nl/)

#### Supplementary Note 2. Psychometric-function analyses

Psychometric functions were fitted to localisation accuracy separately for each participant and expression-by-gaze condition. The main psychometric summary measures were the 75%

localisation threshold, the just-noticeable difference (JND), and the slope parameter. Full fixed-effect statistics are reported in **Supplementary Table S1**. The curve-level accuracy model is reported in **Supplementary Table S2**.

**Supplementary Table S1**

*Repeated-measures analyses of psychometric-function summary measures in Experiment 1.*

| **Metric** | **Effect** | $\boldsymbol{F}$**(df1, df2)** | $\boldsymbol{p}$ | $\eta_p^2$ |
|---|---|---|---|---|
| 75% threshold | Expression | 1.287 (1, 29) | 0.266 | 0.042 |
| 75% threshold | Gaze direction | 4.263 (1, 29) | 0.048 | 0.128 |
| 75% threshold | Expression x Gaze direction | 0.837 (1, 29) | 0.368 | 0.028 |
| JND | Expression | 0.892 (1, 29) | 0.353 | 0.03 |
| JND | Gaze direction | 5.238 (1, 29) | 0.03 | 0.153 |
| JND | Expression x Gaze direction | 0.119 (1, 29) | 0.733 | 0.004 |
| slope beta | Expression | 0.719 (1, 29) | 0.404 | 0.024 |
| slope beta | Gaze duration | 37.49 (1, 29) | < 0.001 | 0.564 |
| slope beta | Expression x Gaze direction | 0.013 (1, 29) | 0.911 | 0 |

*Note.* F statistics are reported as $F$(df1, df2). $\eta_p^2$ = partial eta squared.

**Supplementary Table S2**

*Curve-level localisation accuracy model in Experiment 1.*

| **Effect** | **b** | **SE** | **OR** | **95% CI** | **t(df)** | $\boldsymbol{p}$ |
|---|---|---|---|---|---|---|
| Expression | -0.011 | 0.044 | 0.99 | 0.908 to 1.079 | -0.24 (592) | 0.81 |
| Gaze direction | -0.142 | 0.043 | 0.868 | 0.798 to 0.944 | -3.324 (592) | < 0.001 |
| Exposure duration | 0.995 | 0.069 | 2.703 | 2.362 to 3.094 | 14.46 (592) | < 0.001 |

*Note.* The model was fitted to localisation accuracy across exposure durations. b values are on the logit scale. OR = odds ratio.

#### Supplementary Note 3. Type-1 signal detection analyses

Type-1 signal detection analyses were conducted separately for localisation, expression categorisation, and gaze-direction categorisation. Omnibus repeated-measures analyses are reported in **Supplementary Table S3**. Follow-up tests for significant exposure-dependent effects are reported in **Supplementary Table S4**. Condition-wise tests against zero for sensitivity and response-bias/criterion measures are reported in **Supplementary Table S5**.

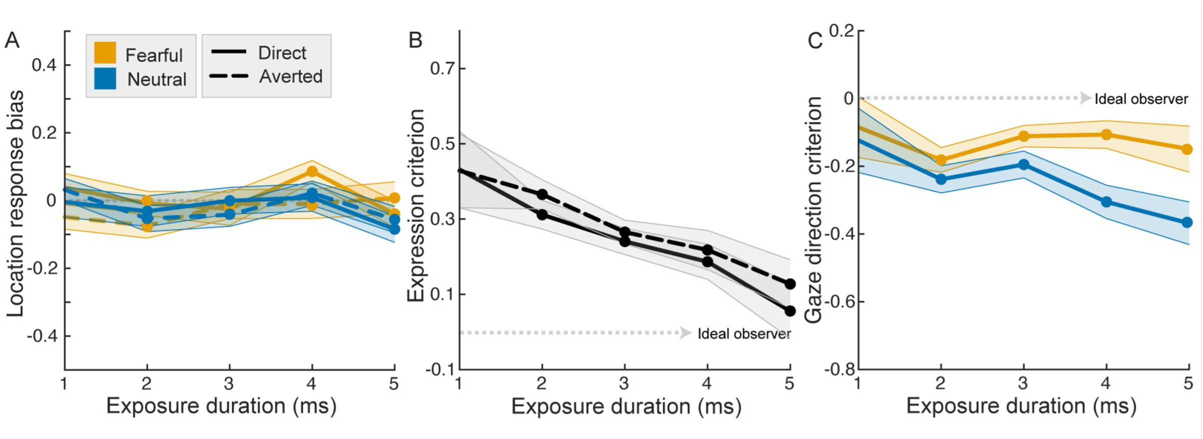


**Supplementary Fig. 2. Response bias and criterion analyses in Experiment 1.** (**A**) Localisation response bias across exposure duration, plotted separately for fearful-direct, fearful-averted, neutral-direct and neutral-averted faces. Values near zero indicate little systematic response tendency in the localisation task. (**B**) Expression criterion across exposure duration, shown separately for direct- and averted-gaze faces. (**C**) Gaze direction criterion across exposure duration, shown separately for fearful and neutral expressions. Shaded regions indicate within-subject SEM. These analyses complement the type-1 sensitivity results by showing response tendencies across tasks and conditions.

#### Supplementary Table S3

*Omnibus type-1 signal detection analyses in Experiment 1.*

| **Task** | **Metric** | **Effect** | ***F*(df1, df2)** | ***p*** | $\eta_p^2$ |
|---|---|---|---|---|---|
| Location | d' | Expression | 0.042 (1, 29) | 0.838 | 0.001 |
| Location | d' | Gaze direction | 25.37 (1, 29) | $< 0.001$ | 0.467 |
| Location | d' | Exposure | 172.9 (4, 116) | $< 0.001$ | 0.856 |
| Location | d' | Expression x Gaze direction | 0.262 (1, 29) | 0.612 | 0.009 |
| Location | d' | Expression x Exposure | 0.68 (4, 116) | 0.565 | 0.023 |
| Location | d' | Gaze direction x Exposure | 17.72 (4, 116) | $< 0.001$ | 0.379 |
| Location | d' | Expression x Gaze direction x Exposure | 0.285 (4, 116) | 0.854 | 0.01 |
| Location | bias | Expression | 0.464 (1, 29) | 0.501 | 0.016 |
| Location | bias | Gaze direction | 1.99 (1, 29) | 0.169 | 0.064 |
| Location | bias | Exposure | 1.487 (4, 116) | 0.23 | 0.049 |
| Location | bias | Expression x Gaze direction | 0.965 (1, 29) | 0.334 | 0.032 |
| Location | bias | Expression x Exposure | 0.639 (4, 116) | 0.603 | 0.022 |
| Location | bias | Gaze direction x Exposure | 0.754 (4, 116) | 0.533 | 0.025 |
| Location | bias | Expression x Gaze direction x Exposure | 1.193 (4, 116) | 0.318 | 0.04 |
| Expression | d' | Gaze direction | 8.389 (1, 29) | 0.007 | 0.224 |
| Expression | d' | Exposure | 36.99 (4, 116) | $< 0.001$ | 0.561 |

| Task | Metric | Effect | ***F*(df1, df2)** | ***p*** | $\eta_p^2$ |
|---|---|---|---|---|---|
| Expression | d' | Gaze direction x Exposure | 1.293 (4, 116) | 0.28 | 0.043 |
| Expression | bias | Gaze direction | 8.655 (1, 29) | 0.006 | 0.23 |
| Expression | bias | Exposure | 4.528 (4, 116) | 0.03 | 0.135 |
| Expression | bias | Gaze direction x Exposure | 0.74 (4, 116) | 0.548 | 0.025 |
| Gaze direction | d' | Expression | 2.8 (1, 29) | 0.105 | 0.088 |
| Gaze direction | d' | Exposure | 22.8 (4, 116) | < 0.001 | 0.44 |
| Gaze direction | d' | Expression x Exposure | 0.046 (4, 116) | 0.993 | 0.002 |
| Gaze direction | bias | Expression | 49.5 (1, 29) | < 0.001 | 0.631 |
| Gaze direction | bias | Exposure | 1.075 (4, 116) | 0.332 | 0.036 |
| Gaze direction | bias | Expression x Exposure | 6.998 (4, 116) | < 0.001 | 0.194 |

*Note.* For the location task, bias indexes left-right response tendency. For expression and gaze direction categorisation tasks, bias corresponds to the response criterion. F statistics are reported as *F*(df1, df2).

**Supplementary Table S4**

*Follow-up tests for type-1 signal detection effects in Experiment 1.*

| Task | Metric | Follow-up | ms | Levels | M diff (SE) | t(df) | *p*Holm | dz | Sig. |
|---|---|---|---|---|---|---|---|---|---|
| Location | d' | Gaze direction within Exposure | 1 | direct - averted | -0.072 (0.044) | -1.641 (29) | 0.223 | -0.3 | No |
| Location | d' | Gaze direction within Exposure | 2 | direct - averted | -0.071 (0.06) | -1.176 (29) | 0.249 | -0.215 | No |
| Location | d' | Gaze direction within Exposure | 3 | direct - averted | 0.045 (0.011) | 4.043 (29) | 0.001 | 0.738 | Yes |
| Location | d' | Gaze direction within Exposure | 4 | direct - averted | 0.339 (0.053) | 6.383 (29) | < 0.001 | 1.165 | Yes |
| Location | d' | Gaze within Exposure | 5 | direct - averted | 0.344 (0.065) | 5.31 (29) | < 0.001 | 0.97 | Yes |
| Gaze direction | bias | Expression within Exposure | 1 | fearful - neutral | 0.039 (0.025) | 1.519 (29) | 0.194 | 0.277 | No |
| Gaze direction | bias | Expression within Exposure | 2 | fearful - neutral | 0.057 (0.033) | 1.716 (29) | 0.194 | 0.313 | No |

| Task | Metric | Follow-up | ms | Levels | M diff (SE) | t(df) | *p*Holm | dz | Sig. |
|---|---|---|---|---|---|---|---|---|---|
| Gaze direction | bias | Expression within Exposure | 3 | fearful - neutral | 0.084 (0.036) | 2.349 (29) | 0.078 | 0.429 | No |
| Gaze direction | bias | Expression within Exposure | 4 | fearful - neutral | 0.199 (0.034) | 5.914 (29) | < 0.001 | 1.08 | Yes |
| Gaze direction | bias | Expression within Exposure | 5 | fearful - neutral | 0.219 (0.036) | 6.112 (29) | < 0.001 | 1.116 | Yes |

*Note.* Mean differences are shown as M diff (SE). Positive values indicate higher values for the first level than the second level. *p*Holm = Holm-corrected *p* value.

**Supplementary Table S5**

*Condition-wise type-1 signal detection tests against zero in Experiment 1.*

| Task | Metric | Condition | ms | M (SE) | t(df) | *p*Holm | dz | Sig. |
|---|---|---|---|---|---|---|---|---|
| Location | d' | fearful-direct | 1 | -0.087 (0.047) | -1.843 (29) | 0.962 | -0.336 | No |
| Location | d' | fearful-direct | 2 | 0.292 (0.054) | 5.44 (29) | < 0.001 | 0.993 | Yes |
| Location | d' | fearful-direct | 3 | 1.159 (0.103) | 11.22 (29) | < 0.001 | 2.048 | Yes |
| Location | d' | fearful-direct | 4 | 1.824 (0.108) | 16.94 (29) | < 0.001 | 3.092 | Yes |
| Location | d' | fearful-direct | 5 | 2.055 (0.136) | 15.16 (29) | < 0.001 | 2.768 | Yes |
| Location | d' | fearful-averted | 1 | 0.023 (0.057) | 0.414 (29) | 0.341 | 0.076 | No |
| Location | d' | fearful-averted | 2 | 0.354 (0.047) | 7.589 (29) | < 0.001 | 1.385 | Yes |
| Location | d' | fearful-averted | 3 | 1.16 (0.121) | 9.576 (29) | < 0.001 | 1.748 | Yes |
| Location | d' | fearful-averted | 4 | 1.463 (0.107) | 13.71 (29) | < 0.001 | 2.502 | Yes |
| Location | d' | fearful-averted | 5 | 1.725 (0.103) | 16.75 (29) | < 0.001 | 3.058 | Yes |
| Location | d' | neutral-direct | 1 | -0.037 (0.049) | -0.756 (29) | 0.772 | -0.138 | No |
| Location | d' | neutral-direct | 2 | 0.288 (0.045) | 6.444 (29) | < 0.001 | 1.177 | Yes |
| Location | d' | neutral-direct | 3 | 1.127 (0.105) | 10.78 (29) | < 0.001 | 1.968 | Yes |
| Location | d' | neutral-direct | 4 | 1.835 (0.113) | 16.28 (29) | < 0.001 | 2.973 | Yes |
| Location | d' | neutral-direct | 5 | 2.074 (0.138) | 15.08 (29) | < 0.001 | 2.753 | Yes |
| Location | d' | neutral-averted | 1 | -0.002 (0.064) | -0.038 (29) | 0.515 | -0.007 | No |

| Task | Metric | Condition | ms | M (SE) | t(df) | *p*Holm | dz | Sig. |
|---|---|---|---|---|---|---|---|---|
| Location | d' | neutral-averted | 2 | 0.367 (0.067) | 5.461 (29) | < 0.001 | 0.997 | Yes |
| Location | d' | neutral-averted | 3 | 1.037 (0.086) | 12.13 (29) | < 0.001 | 2.214 | Yes |
| Location | d' | neutral-averted | 4 | 1.517 (0.118) | 12.81 (29) | < 0.001 | 2.339 | Yes |
| Location | d' | neutral-averted | 5 | 1.716 (0.108) | 15.94 (29) | < 0.001 | 2.91 | Yes |
| Location | bias | fearful-direct | 1 | 0.038 (0.029) | 1.337 (29) | 0.766 | 0.244 | No |
| Location | bias | fearful-direct | 2 | -0.009 (0.051) | -0.171 (29) | 1 | -0.031 | No |
| Location | bias | fearful-direct | 3 | -0.024 (0.058) | -0.415 (29) | 1 | -0.076 | No |
| Location | bias | fearful-direct | 4 | 0.086 (0.032) | 2.686 (29) | 0.059 | 0.49 | No |
| Location | bias | fearful-direct | 5 | -0.04 (0.035) | -1.159 (29) | 0.768 | -0.212 | No |
| Location | bias | fearful-averted | 1 | -0.049 (0.025) | -1.923 (29) | 0.322 | -0.351 | No |
| Location | bias | fearful-averted | 2 | -0.074 (0.05) | -1.478 (29) | 0.601 | -0.27 | No |
| Location | bias | fearful-averted | 3 | -0.01 (0.053) | -0.195 (29) | 1 | -0.036 | No |
| Location | bias | fearful-averted | 4 | -0.01 (0.051) | -0.194 (29) | 1 | -0.035 | No |
| Location | bias | fearful-averted | 5 | 0.008 (0.052) | 0.153 (29) | 1 | 0.028 | No |
| Location | bias | neutral-direct | 1 | -0.005 (0.03) | -0.153 (29) | 1 | -0.028 | No |
| Location | bias | neutral-direct | 2 | -0.031 (0.057) | -0.547 (29) | 1 | -0.1 | No |
| Location | bias | neutral-direct | 3 | -0.001 (0.054) | -0.009 (29) | 1 | -0.002 | No |
| Location | bias | neutral-direct | 4 | 0.009 (0.043) | 0.21 (29) | 1 | 0.038 | No |
| Location | bias | neutral-direct | 5 | -0.084 (0.043) | -1.969 (29) | 0.293 | -0.36 | No |
| Location | bias | neutral-averted | 1 | 0.032 (0.029) | 1.116 (29) | 1 | 0.204 | No |
| Location | bias | neutral-averted | 2 | -0.053 (0.044) | -1.205 (29) | 1 | -0.22 | No |
| Location | bias | neutral-averted | 3 | -0.042 (0.05) | -0.831 (29) | 1 | -0.152 | No |
| Location | bias | neutral-averted | 4 | 0.022 (0.049) | 0.449 (29) | 1 | 0.082 | No |
| Location | bias | neutral-averted | 5 | -0.056 (0.044) | -1.273 (29) | 1 | -0.232 | No |
| Expression | d' | direct | 1 | 0.032 (0.053) | 0.603 (29) | 0.551 | 0.11 | No |
| Expression | d' | direct | 2 | -0.077 (0.065) | -1.183 (29) | 0.877 | -0.216 | No |

| Task | Metric | Condition | ms | M (SE) | t(df) | *p*Holm | dz | Sig. |
|---|---|---|---|---|---|---|---|---|
| Expression | d' | direct | 3 | 0.103 (0.046) | 2.234 (29) | 0.05 | 0.408 | No |
| Expression | d' | direct | 4 | 0.333 (0.055) | 6.017 (29) | < 0.001 | 1.098 | Yes |
| Expression | d' | direct | 5 | 0.663 (0.079) | 8.342 (29) | < 0.001 | 1.523 | Yes |
| Expression | d' | averted | 1 | 0.044 (0.055) | 0.812 (29) | 0.463 | 0.148 | No |
| Expression | d' | averted | 2 | -0.15 (0.049) | -3.061 (29) | 1 | -0.559 | No |
| Expression | d' | averted | 3 | 0.06 (0.058) | 1.037 (29) | 0.463 | 0.189 | No |
| Expression | d' | averted | 4 | 0.163 (0.039) | 4.216 (29) | < 0.001 | 0.77 | Yes |
| Expression | d' | averted | 5 | 0.497 (0.061) | 8.162 (29) | < 0.001 | 1.49 | Yes |
| Expression | bias | direct | 1 | 0.432 (0.137) | 3.157 (29) | 0.007 | 0.576 | Yes |
| Expression | bias | direct | 2 | 0.312 (0.083) | 3.769 (29) | 0.003 | 0.688 | Yes |
| Expression | bias | direct | 3 | 0.241 (0.062) | 3.887 (29) | 0.003 | 0.71 | Yes |
| Expression | bias | direct | 4 | 0.187 (0.048) | 3.922 (29) | 0.003 | 0.716 | Yes |
| Expression | bias | direct | 5 | 0.056 (0.044) | 1.265 (29) | 0.216 | 0.231 | No |
| Expression | bias | averted | 1 | 0.429 (0.136) | 3.163 (29) | 0.007 | 0.577 | Yes |
| Expression | bias | averted | 2 | 0.366 (0.086) | 4.253 (29) | < 0.001 | 0.777 | Yes |
| Expression | bias | averted | 3 | 0.265 (0.057) | 4.658 (29) | < 0.001 | 0.85 | Yes |
| Expression | bias | averted | 4 | 0.218 (0.054) | 4.042 (29) | 0.001 | 0.738 | Yes |
| Expression | bias | averted | 5 | 0.128 (0.06) | 2.126 (29) | 0.042 | 0.388 | Yes |
| Gaze direction | d' | fearful | 1 | 0.016 (0.042) | 0.385 (29) | 1 | 0.07 | No |
| Gaze direction | d' | fearful | 2 | 0.001 (0.047) | 0.029 (29) | 1 | 0.005 | No |
| Gaze direction | d' | fearful | 3 | 0.002 (0.045) | 0.04 (29) | 1 | 0.007 | No |
| Gaze direction | d' | fearful | 4 | 0.15 (0.055) | 2.74 (29) | 0.021 | 0.5 | Yes |
| Gaze direction | d' | fearful | 5 | 0.386 (0.047) | 8.2 (29) | < 0.001 | 1.497 | Yes |
| Gaze direction | d' | neutral | 1 | 0.048 (0.056) | 0.846 (29) | 0.286 | 0.154 | No |
| Gaze direction | d' | neutral | 2 | 0.046 (0.042) | 1.087 (29) | 0.286 | 0.198 | No |
| Gaze direction | d' | neutral | 3 | 0.073 (0.051) | 1.411 (29) | 0.253 | 0.258 | No |

| Task | Metric | Condition | ms | M (SE) | t(df) | *p*Holm | dz | Sig. |
|---|---|---|---|---|---|---|---|---|
| Gaze direction | d' | neutral | 4 | 0.208 (0.051) | 4.062 (29) | < 0.001 | 0.742 | Yes |
| Gaze direction | d' | neutral | 5 | 0.427 (0.051) | 8.306 (29) | < 0.001 | 1.516 | Yes |
| Gaze direction | bias | fearful | 1 | -0.085 (0.133) | -0.642 (29) | 0.526 | -0.117 | No |
| Gaze direction | bias | fearful | 2 | -0.181 (0.091) | -1.988 (29) | 0.225 | -0.363 | No |
| Gaze direction | bias | fearful | 3 | -0.111 (0.076) | -1.464 (29) | 0.308 | -0.267 | No |
| Gaze direction | bias | fearful | 4 | -0.107 (0.063) | -1.702 (29) | 0.298 | -0.311 | No |
| Gaze direction | bias | fearful | 5 | -0.149 (0.04) | -3.718 (29) | 0.004 | -0.679 | Yes |
| Gaze direction | bias | neutral | 1 | -0.124 (0.133) | -0.932 (29) | 0.359 | -0.17 | No |
| Gaze direction | bias | neutral | 2 | -0.239 (0.089) | -2.688 (29) | 0.035 | -0.491 | Yes |
| Gaze direction | bias | neutral | 3 | -0.195 (0.076) | -2.566 (29) | 0.035 | -0.469 | Yes |
| Gaze direction | bias | neutral | 4 | -0.305 (0.063) | -4.81 (29) | < 0.001 | -0.878 | Yes |
| Gaze direction | bias | neutral | 5 | -0.368 (0.05) | -7.36 (29) | < 0.001 | -1.344 | Yes |

*Note.* For sensitivity measures, positive values indicate above-chance performance. For bias/criterion measures, values different from zero indicate response bias. *p*Holm = Holm-corrected *p* value.

#### Supplementary Note 4. Additional metacognitive analyses

This note reports the additional type-2 signal detection and hierarchical metacognitive-efficiency analyses for Experiment 1. Meta-*d′* quantified how well perceptual awareness ratings tracked localisation performance, whereas M-ratio quantified metacognitive efficiency relative to type-1 sensitivity.

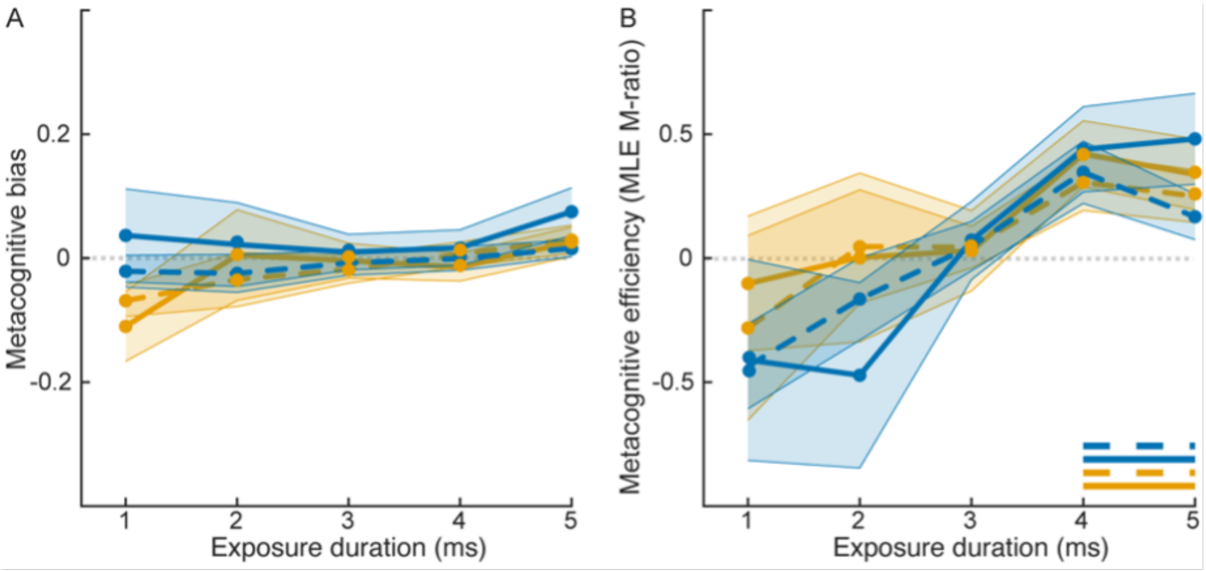


**Supplementary Fig. 3. Metacognitive bias and metacognitive efficiency (MLE) in Experiment 1.** (**A**) Metacognitive bias derived from the type-2 signal detection analysis of perceptual awareness ratings in the localisation task, shown across exposure durations and stimulus conditions. (**B**) Metacognitive efficiency, quantified as maximum-likelihood M-ratio, across exposure durations and stimulus conditions. Values indicate how efficiently subjective awareness tracked the perceptual evidence available for type-1 localisation performance. Shaded regions indicate within-subject SEM. These analyses complement the meta-$d'$ and hierarchical M-ratio results reported in the main text.

#### Supplementary Table S6

*Omnibus type-2 signal detection analyses in Experiment 1.*

| Metric | Effect | F(df1, df2) | $p$ | $\eta_p^2$ |
|---|---|---|---|---|
| meta-d' | Expression | 0.408 (1, 27) | 0.528 | 0.015 |
| meta-d' | Gaze direction | 18.23 (1, 27) | < 0.001 | 0.403 |
| meta-d' | Exposure | 25.26 (4, 108) | < 0.001 | 0.483 |
| meta-d' | Expression x Gaze direction | 0.044 (1, 27) | 0.835 | 0.002 |
| meta-d' | Expression x Exposure | 0.144 (4, 108) | 0.933 | 0.005 |
| meta-d' | Gaze direction x Exposure | 6.43 (4, 108) | < 0.001 | 0.192 |
| meta-d' | Expression x Gaze direction x Exposure | 0.233 (4, 108) | 0.867 | 0.009 |
| meta-bias | Expression | 3.49 (1, 27) | 0.073 | 0.114 |
| meta-bias | Gaze direction | 1.714 (1, 27) | 0.201 | 0.06 |
| meta-bias | Exposure | 1.858 (4, 108) | 0.162 | 0.064 |
| meta-bias | Expression x Gaze direction | 1.747 (1, 27) | 0.197 | 0.061 |
| meta-bias | Expression x Exposure | 1.11 (4, 108) | 0.346 | 0.04 |
| meta-bias | Gaze direction x Exposure | 0.308 (4, 108) | 0.72 | 0.011 |
| meta-bias | Expression x Gaze direction x Exposure | 0.23 (4, 108) | 0.825 | 0.008 |
| M-ratio | Expression | 0.618 (1, 27) | 0.439 | 0.022 |
| M-ratio | Gaze direction | 0.441 (1, 27) | 0.512 | 0.016 |
| M-ratio | Exposure | 7.902 (4, 108) | < 0.001 | 0.226 |

| Metric | Effect | F(df1, df2) | *p* | $\eta_p^2$ |
|---|---|---|---|---|
| M-ratio | Expression x Gaze direction | 0.006 (1, 27) | 0.94 | 0 |
| M-ratio | Expression x Exposure | 0.361 (4, 108) | 0.667 | 0.013 |
| M-ratio | Gaze direction x Exposure | 0.262 (4, 108) | 0.78 | 0.01 |
| M-ratio | Expression x Gaze direction x Exposure | 0.23 (4, 108) | 0.812 | 0.008 |

*Note.* F statistics are reported as $F$(df1, df2). $\eta_p^2$ = partial eta squared.

**Supplementary Table S7**

*Follow-up tests for meta-d′ gaze effects across exposure duration.*

| Metric | Follow-up | ms | M diff (SE) | t(df) | *p*Holm | dz | Sig. |
|---|---|---|---|---|---|---|---|
| meta-d' | Gaze direction within Exposure | 1 | -0.006 (0.063) | -0.091 (29) | 0.928 | -0.017 | No |
| meta-d' | Gaze direction within Exposure | 2 | 0.098 (0.068) | 1.437 (29) | 0.323 | 0.262 | No |
| meta-d' | Gaze direction within Exposure | 3 | -0.092 (0.049) | -1.864 (29) | 0.217 | -0.34 | No |
| meta-d' | Gaze direction within Exposure | 4 | 0.382 (0.089) | 4.313 (29) | < 0.001 | 0.787 | Yes |
| meta-d' | Gaze direction within Exposure | 5 | 0.368 (0.115) | 3.196 (29) | 0.013 | 0.584 | Yes |

*Note.* Mean differences are shown as M diff (SE). Positive values indicate higher meta-*d′* for direct gaze than averted gaze. *p*Holm = Holm-corrected p value.

**Supplementary Table S8**

*Condition-wise type-2 signal detection tests against zero in Experiment 1.*

| Metric | Condition | ms | M (SE) | t(df) | *p*Holm | dz | Sig. |
|---|---|---|---|---|---|---|---|
| meta-d' | fearful-direct | 1 | 0.015 (0.058) | 0.251 (28) | 1 | 0.047 | No |
| meta-d' | fearful-direct | 2 | 0.022 (0.062) | 0.345 (29) | 1 | 0.063 | No |
| meta-d' | fearful-direct | 3 | -0.093 (0.076) | -1.227 (29) | 1 | -0.224 | No |
| meta-d' | fearful-direct | 4 | 0.663 (0.118) | 5.621 (29) | < 0.001 | 1.026 | Yes |
| meta-d' | fearful-direct | 5 | 0.673 (0.128) | 5.276 (29) | < 0.001 | 0.963 | Yes |

| Metric | Condition | ms | M (SE) | t(df) | *p*Holm | dz | Sig. |
|---|---|---|---|---|---|---|---|
| meta-d' | fearful-averted | 1 | -0.008 (0.063) | -0.12 (29) | 1 | -0.022 | No |
| meta-d' | fearful-averted | 2 | -0.061 (0.056) | -1.089 (28) | 1 | -0.202 | No |
| meta-d' | fearful-averted | 3 | 0.03 (0.059) | 0.517 (29) | 0.913 | 0.094 | No |
| meta-d' | fearful-averted | 4 | 0.227 (0.122) | 1.871 (29) | 0.143 | 0.342 | No |
| meta-d' | fearful-averted | 5 | 0.345 (0.14) | 2.473 (29) | 0.049 | 0.452 | Yes |
| meta-d' | neutral-direct | 1 | -0.018 (0.058) | -0.304 (29) | 1 | -0.056 | No |
| meta-d' | neutral-direct | 2 | 0.015 (0.055) | 0.282 (29) | 1 | 0.052 | No |
| meta-d' | neutral-direct | 3 | -0.021 (0.08) | -0.259 (29) | 1 | -0.047 | No |
| meta-d' | neutral-direct | 4 | 0.672 (0.124) | 5.433 (29) | < 0.001 | 0.992 | Yes |
| meta-d' | neutral-direct | 5 | 0.733 (0.129) | 5.686 (29) | < 0.001 | 1.038 | Yes |
| meta-d' | neutral-averted | 1 | 0.018 (0.057) | 0.309 (29) | 0.903 | 0.056 | No |
| meta-d' | neutral-averted | 2 | -0.081 (0.078) | -1.049 (29) | 0.903 | -0.191 | No |
| meta-d' | neutral-averted | 3 | 0.04 (0.075) | 0.527 (29) | 0.903 | 0.096 | No |
| meta-d' | neutral-averted | 4 | 0.342 (0.112) | 3.039 (29) | 0.012 | 0.555 | Yes |
| meta-d' | neutral-averted | 5 | 0.325 (0.106) | 3.067 (29) | 0.012 | 0.56 | Yes |
| meta-bias | fearful-direct | 1 | -0.11 (0.053) | -2.085 (28) | 0.232 | -0.387 | No |
| meta-bias | fearful-direct | 2 | 0.005 (0.078) | 0.064 (29) | 1 | 0.012 | No |
| meta-bias | fearful-direct | 3 | -0.004 (0.021) | -0.188 (29) | 1 | -0.034 | No |
| meta-bias | fearful-direct | 4 | -0.015 (0.016) | -0.906 (29) | 1 | -0.165 | No |
| meta-bias | fearful-direct | 5 | 0.026 (0.015) | 1.763 (29) | 0.353 | 0.322 | No |
| meta-bias | fearful-averted | 1 | -0.069 (0.037) | -1.861 (29) | 0.364 | -0.34 | No |
| meta-bias | fearful-averted | 2 | -0.034 (0.053) | -0.648 (28) | 1 | -0.12 | No |
| meta-bias | fearful-averted | 3 | -0.018 (0.013) | -1.402 (29) | 0.514 | -0.256 | No |
| meta-bias | fearful-averted | 4 | 0.006 (0.017) | 0.361 (29) | 1 | 0.066 | No |
| meta-bias | fearful-averted | 5 | 0.031 (0.018) | 1.706 (29) | 0.395 | 0.311 | No |
| meta-bias | neutral-direct | 1 | 0.037 (0.077) | 0.476 (29) | 1 | 0.087 | No |

| Metric | Condition | ms | M (SE) | t(df) | $p$Holm | dz | Sig. |
|---|---|---|---|---|---|---|---|
| meta-bias | neutral-direct | 2 | 0.022 (0.07) | 0.315 (29) | 1 | 0.058 | No |
| meta-bias | neutral-direct | 3 | 0.01 (0.019) | 0.517 (29) | 1 | 0.094 | No |
| meta-bias | neutral-direct | 4 | 0.017 (0.022) | 0.782 (29) | 1 | 0.143 | No |
| meta-bias | neutral-direct | 5 | 0.075 (0.033) | 2.272 (29) | 0.154 | 0.415 | No |
| meta-bias | neutral-averted | 1 | -0.021 (0.026) | -0.823 (29) | 1 | -0.15 | No |
| meta-bias | neutral-averted | 2 | -0.025 (0.032) | -0.769 (29) | 1 | -0.14 | No |
| meta-bias | neutral-averted | 3 | -0.008 (0.017) | -0.445 (29) | 1 | -0.081 | No |
| meta-bias | neutral-averted | 4 | -0.001 (0.018) | -0.035 (29) | 1 | -0.006 | No |
| meta-bias | neutral-averted | 5 | 0.016 (0.012) | 1.3 (29) | 1 | 0.237 | No |
| M-ratio | fearful-direct | 1 | -0.102 (0.323) | -0.316 (28) | 1 | -0.059 | No |
| M-ratio | fearful-direct | 2 | 0.003 (0.375) | 0.008 (29) | 1 | 0.001 | No |
| M-ratio | fearful-direct | 3 | 0.03 (0.146) | 0.206 (29) | 1 | 0.038 | No |
| M-ratio | fearful-direct | 4 | 0.421 (0.079) | 5.33 (29) | < 0.001 | 0.973 | Yes |
| M-ratio | fearful-direct | 5 | 0.338 (0.059) | 5.76 (29) | < 0.001 | 1.052 | Yes |
| M-ratio | fearful-averted | 1 | -0.281 (0.378) | -0.744 (29) | 0.802 | -0.136 | No |
| M-ratio | fearful-averted | 2 | 0.047 (0.185) | 0.253 (28) | 0.802 | 0.047 | No |
| M-ratio | fearful-averted | 3 | 0.045 (0.065) | 0.69 (29) | 0.744 | 0.126 | No |
| M-ratio | fearful-averted | 4 | 0.306 (0.107) | 2.844 (29) | 0.016 | 0.519 | Yes |
| M-ratio | fearful-averted | 5 | 0.25 (0.079) | 3.184 (29) | 0.009 | 0.581 | Yes |
| M-ratio | neutral-direct | 1 | -0.41 (0.456) | -0.899 (29) | 1 | -0.164 | No |
| M-ratio | neutral-direct | 2 | -0.472 (0.402) | -1.174 (29) | 1 | -0.214 | No |
| M-ratio | neutral-direct | 3 | 0.071 (0.094) | 0.761 (29) | 0.68 | 0.139 | No |
| M-ratio | neutral-direct | 4 | 0.439 (0.085) | 5.155 (29) | < 0.001 | 0.941 | Yes |
| M-ratio | neutral-direct | 5 | 0.481 (0.124) | 3.891 (29) | 0.001 | 0.71 | Yes |
| M-ratio | neutral-averted | 1 | -0.436 (0.173) | -2.513 (29) | 1 | -0.459 | No |
| M-ratio | neutral-averted | 2 | -0.165 (0.179) | -0.92 (29) | 1 | -0.168 | No |

| Metric | Condition | ms | M (SE) | t(df) | *p*Holm | dz | Sig. |
|---|---|---|---|---|---|---|---|
| M-ratio | neutral-averted | 3 | 0.049 (0.079) | 0.615 (29) | 0.815 | 0.112 | No |
| M-ratio | neutral-averted | 4 | 0.346 (0.11) | 3.157 (29) | 0.009 | 0.576 | Yes |
| M-ratio | neutral-averted | 5 | 0.167 (0.07) | 2.402 (29) | 0.046 | 0.439 | Yes |

*Note.* Condition-wise tests are reported for meta-*d*′, M-ratio, and metacognitive bias. *p*Holm = Holm-corrected *p* value.

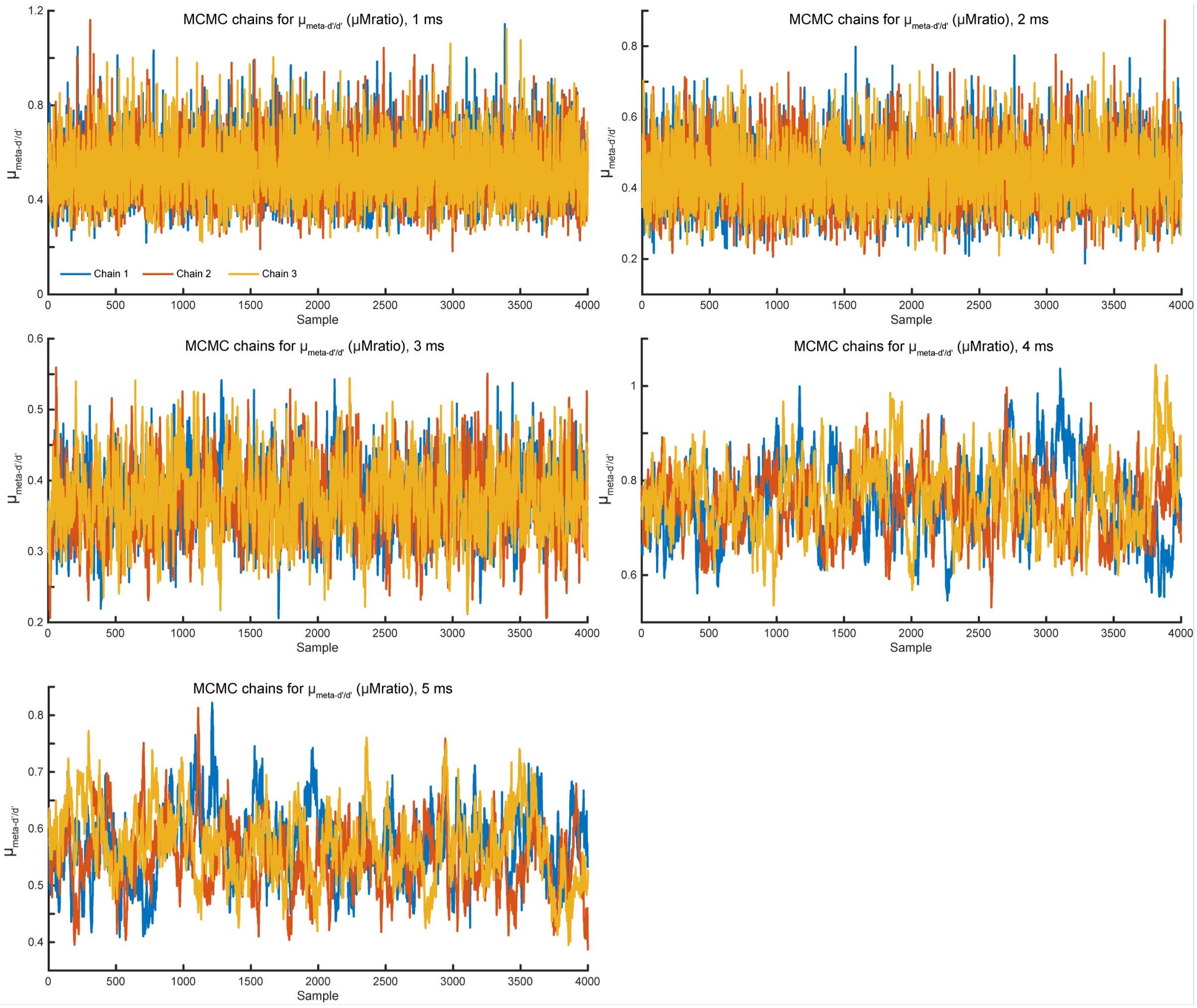


**Supplementary Fig. 4. Hierarchical metacognitive model convergence diagnostics.** MCMC trace plots for the hierarchical meta-*d*′ analyses estimating group-level metacognitive efficiency across exposure durations. Panels show the posterior sampling chains for the group-level M-ratio parameter at 1, 2, 3, 4 and 5 ms. Independent chains are shown in different colours. These diagnostic plots were inspected to assess chain mixing and convergence of the hierarchical metacognitive-efficiency model.

#### Supplementary Table S9

*Hierarchical Bayesian M-ratio contrasts between direct and averted gaze.*

| ms | Direct | Averted | $\beta$ | 95% CrI | $P$(direct > averted) | Decision | DIC |
|---|---|---|---|---|---|---|---|
| 1 | 0.672 | 0.562 | -0.258 | -0.820 to 0.293 | 0.821 | CrI includes zero | 1493.6 |
| 2 | 0.471 | 0.328 | -0.389 | -0.886 to 0.098 | 0.94 | CrI includes zero | 1896.4 |
| 3 | 0.37 | 0.393 | 0.006 | -0.368 to 0.357 | 0.481 | CrI includes zero | 2151.3 |
| 4 | 0.77 | 0.568 | -0.321 | -0.616 to -0.035 | 0.986 | direct > averted | 2267.9 |
| 5 | 0.596 | 0.356 | -0.502 | -0.857 to -0.165 | 0.998 | direct > averted | 2258.5 |

*Note.* $\beta$ is the group-level contrast on the log scale. CrI = credible interval; DIC = deviance information criterion.

#### Supplementary Note 5. Information-theoretic analyses

This note reports the full information-theoretic statistics for Experiment 1. Mutual information estimates quantified the amount of stimulus information carried by behavioural responses. PAS co-information quantified whether subjective visibility ratings overlapped with, or modulated, the information shared between stimulus location and localisation responses. Bias-corrected estimates can fall slightly below zero and should be interpreted as no evidence for information above the permutation baseline.

##### Supplementary Table S10

*Condition-wise mutual information tests against zero in Experiment 1.*

| Task | Condition | ms | M (SE) | t(df) | $p$Holm | dz | Sig. |
|---|---|---|---|---|---|---|---|
| Expression | averted | 1 | -0.001 (0.002) | -0.603 (29) | 1 | -0.11 | No |
| Expression | averted | 2 | 0.001 (0.002) | 0.665 (29) | 1 | 0.121 | No |
| Expression | averted | 3 | 0.003 (0.002) | 1.608 (29) | 0.949 | 0.294 | No |
| Expression | averted | 4 | -0 (0.002) | -0.143 (29) | 1 | -0.026 | No |
| Expression | averted | 5 | 0.03 (0.006) | 4.753 (29) | < 0.001 | 0.868 | Yes |
| Expression | direct | 1 | -0.001 (0.001) | -0.664 (29) | 1 | -0.121 | No |
| Expression | direct | 2 | 0.005 (0.003) | 1.831 (29) | 0.664 | 0.334 | No |
| Expression | direct | 3 | -0 (0.002) | -0.01 (29) | 1 | -0.002 | No |
| Expression | direct | 4 | 0.014 (0.004) | 3.399 (29) | 0.022 | 0.621 | Yes |

| **Task** | **Condition** | **ms** | **M (SE)** | **t(df)** | ***p*Holm** | **dz** | **Sig.** |
|---|---|---|---|---|---|---|---|
| Expression | direct | 5 | 0.058 (0.011) | 5.176 (29) | < 0.001 | 0.945 | Yes |
| Gaze direction | fearful | 1 | -0.002 (0.002) | -1.251 (28) | 1 | -0.232 | No |
| Gaze direction | fearful | 2 | -0.002 (0.001) | -1.135 (29) | 1 | -0.207 | No |
| Gaze direction | fearful | 3 | -0.002 (0.001) | -1.92 (29) | 1 | -0.35 | No |
| Gaze direction | fearful | 4 | 0.004 (0.003) | 1.438 (29) | 1 | 0.263 | No |
| Gaze direction | fearful | 5 | 0.015 (0.005) | 3.245 (29) | 0.031 | 0.593 | Yes |
| Gaze direction | neutral | 1 | 0.001 (0.002) | 0.545 (28) | 1 | 0.101 | No |
| Gaze direction | neutral | 2 | -0.002 (0.001) | -1.338 (29) | 1 | -0.244 | No |
| Gaze direction | neutral | 3 | 0.001 (0.002) | 0.637 (29) | 1 | 0.116 | No |
| Gaze direction | neutral | 4 | 0.004 (0.002) | 1.855 (29) | 0.664 | 0.339 | No |
| Gaze direction | neutral | 5 | 0.017 (0.004) | 3.869 (29) | 0.007 | 0.706 | Yes |
| Location | fearful-averted | 1 | 0.004 (0.004) | 0.932 (29) | 1 | 0.17 | No |
| Location | fearful-averted | 2 | 0.025 (0.007) | 3.746 (29) | 0.009 | 0.684 | Yes |
| Location | fearful-averted | 3 | 0.263 (0.044) | 5.929 (29) | < 0.001 | 1.083 | Yes |
| Location | fearful-averted | 4 | 0.371 (0.039) | 9.538 (29) | < 0.001 | 1.741 | Yes |
| Location | fearful-averted | 5 | 0.473 (0.04) | 11.85 (29) | < 0.001 | 2.163 | Yes |
| Location | fearful-direct | 1 | -0.001 (0.005) | -0.174 (29) | 1 | -0.032 | No |
| Location | fearful-direct | 2 | 0.021 (0.006) | 3.424 (29) | 0.021 | 0.625 | Yes |
| Location | fearful-direct | 3 | 0.261 (0.035) | 7.397 (29) | < 0.001 | 1.35 | Yes |
| Location | fearful-direct | 4 | 0.514 (0.044) | 11.78 (29) | < 0.001 | 2.15 | Yes |
| Location | fearful-direct | 5 | 0.61 (0.054) | 11.19 (29) | < 0.001 | 2.043 | Yes |
| Location | neutral-averted | 1 | 0.011 (0.004) | 2.883 (29) | 0.074 | 0.526 | No |
| Location | neutral-averted | 2 | 0.041 (0.007) | 5.725 (29) | < 0.001 | 1.045 | Yes |
| Location | neutral-averted | 3 | 0.214 (0.031) | 6.983 (29) | < 0.001 | 1.275 | Yes |
| Location | neutral-averted | 4 | 0.4 (0.045) | 8.875 (29) | < 0.001 | 1.62 | Yes |
| Location | neutral-averted | 5 | 0.471 (0.041) | 11.57 (29) | < 0.001 | 2.112 | Yes |

| Task | Condition | ms | M (SE) | t(df) | *p*Holm | dz | Sig. |
|---|---|---|---|---|---|---|---|
| Location | neutral-direct | 1 | -0 (0.003) | -0.099 (29) | 1 | -0.018 | No |
| Location | neutral-direct | 2 | 0.014 (0.005) | 2.846 (29) | 0.076 | 0.52 | No |
| Location | neutral-direct | 3 | 0.253 (0.037) | 6.749 (29) | < 0.001 | 1.232 | Yes |
| Location | neutral-direct | 4 | 0.519 (0.045) | 11.44 (29) | < 0.001 | 2.089 | Yes |
| Location | neutral-direct | 5 | 0.619 (0.054) | 11.41 (29) | < 0.001 | 2.083 | Yes |

*Note.* Mutual information values are bias-corrected and expressed in bits. $p_{\text{Holm}}$ = Holm-corrected p value.

**Supplementary Table S11**

*Planned information-theoretic contrasts in Experiment 1.*

| Contrast | Condition | ms | M (SE) | t(df) | *p*Holm | dz | Sig. |
|---|---|---|---|---|---|---|---|
| Location: direct-averted | fearful | 1 | -0.005 (0.006) | -0.831 (29) | 1 | -0.152 | No |
| Location: direct-averted | fearful | 2 | -0.004 (0.009) | -0.473 (29) | 1 | -0.086 | No |
| Location: direct-averted | fearful | 3 | -0.002 (0.024) | -0.101 (29) | 1 | -0.018 | No |
| Location: direct-averted | fearful | 4 | 0.143 (0.037) | 3.896 (29) | 0.005 | 0.711 | Yes |
| Location: direct-averted | fearful | 5 | 0.137 (0.033) | 4.194 (29) | 0.002 | 0.766 | Yes |
| Location: direct-averted | neutral | 1 | -0.011 (0.005) | -2.308 (29) | 1 | -0.421 | No |
| Location: direct-averted | neutral | 2 | -0.027 (0.008) | -3.439 (29) | 1 | -0.628 | No |
| Location: direct-averted | neutral | 3 | 0.038 (0.023) | 1.657 (29) | 0.758 | 0.302 | No |
| Location: direct-averted | neutral | 4 | 0.119 (0.031) | 3.892 (29) | 0.005 | 0.711 | Yes |
| Location: direct-averted | neutral | 5 | 0.148 (0.034) | 4.374 (29) | 0.001 | 0.799 | Yes |
| Expression: direct-averted | direct-averted | 1 | 0 (0.002) | 0.029 (29) | 1 | 0.005 | No |

| Contrast | Condition | ms | M (SE) | t(df) | *p*Holm | dz | Sig. |
|---|---|---|---|---|---|---|---|
| Expression: direct-averted | direct-averted | 2 | 0.004 (0.003) | 1.351 (29) | 1 | 0.247 | No |
| Expression: direct-averted | direct-averted | 3 | -0.003 (0.003) | -1.138 (29) | 1 | -0.208 | No |
| Expression: direct-averted | direct-averted | 4 | 0.014 (0.004) | 3.317 (29) | 0.02 | 0.606 | Yes |
| Expression: direct-averted | direct-averted | 5 | 0.027 (0.01) | 2.785 (29) | 0.07 | 0.508 | No |
| Gaze direction: fearful-neutral | fearful-neutral | 1 | -0.004 (0.003) | -1.25 (28) | 1 | -0.232 | No |
| Gaze direction: fearful-neutral | fearful-neutral | 2 | 0 (0.002) | 0.107 (29) | 1 | 0.02 | No |
| Gaze direction: fearful-neutral | fearful-neutral | 3 | -0.003 (0.002) | -1.503 (29) | 1 | -0.275 | No |
| Gaze direction: fearful-neutral | fearful-neutral | 4 | -0 (0.003) | -0.113 (29) | 1 | -0.021 | No |
| Gaze direction: fearful-neutral | fearful-neutral | 5 | -0.002 (0.007) | -0.299 (29) | 1 | -0.055 | No |

*Note.* Contrasts were computed at the participant level before group-level tests. $p_{\text{Holm}}$ = Holm-corrected p value.

**Supplementary Table S12**

Direct minus averted localisation mutual information contrasts collapsed across emotional expression in Experiment 1.

| Contrast | Condition | ms | M diff (SE) | t(df) | $p_{\text{Holm}}$ | dz | Sig. |
|---|---|---|---|---|---|---|---|
| Location: direct minus averted | collapsed across expression | 1 | -0.008 (0.004) | -1.886 (29) | 1.000 | -0.344 | No |
| Location: direct minus averted | collapsed across expression | 2 | -0.016 (0.006) | -2.458 (29) | 1.000 | -0.449 | No |

| Contrast | Condition | ms | M diff (SE) | t(df) | $p_{\text{Holm}}$ | dz | Sig. |
|---|---|---|---|---|---|---|---|
| Location: direct minus averted | collapsed across expression | 3 | 0.018 (0.005) | 3.832 (29) | < 0.001 | 0.700 | Yes |
| Location: direct minus averted | collapsed across expression | 4 | 0.131 (0.020) | 6.449 (29) | < 0.001 | 1.177 | Yes |
| Location: direct minus averted | collapsed across expression | 5 | 0.143 (0.027) | 5.322 (29) | < 0.001 | 0.972 | Yes |

*Note.* Contrasts were computed at the participant level by averaging the direct minus averted localisation mutual information difference across fearful and neutral faces, separately for each exposure duration. Positive values indicate greater localisation information for direct gaze than averted gaze. $p_{\text{Holm}}$ = Holm corrected p value across the five exposure durations. dz = Cohen's dz.

**Supplementary Table S13**

*PAS co-information tests for localisation in Experiment 1.*

| Condition | ms | M (SE) | t(df) | *p*Holm | dz | Sig. |
|---|---|---|---|---|---|---|
| fearful-averted | 1 | 0.003 (0.005) | 0.585 (29) | 1 | 0.107 | No |
| fearful-averted | 2 | 0.005 (0.004) | 1.26 (29) | 1 | 0.23 | No |
| fearful-averted | 3 | -0.047 (0.014) | -3.282 (29) | 0.1 | -0.599 | No |
| fearful-averted | 4 | 0.017 (0.014) | 1.239 (29) | 1 | 0.226 | No |
| fearful-averted | 5 | 0.064 (0.022) | 2.934 (29) | 0.233 | 0.536 | No |
| fearful-direct | 1 | 0.003 (0.006) | 0.53 (28) | 1 | 0.099 | No |
| fearful-direct | 2 | 0.003 (0.004) | 0.732 (29) | 1 | 0.134 | No |
| fearful-direct | 3 | 0.001 (0.009) | 0.156 (29) | 1 | 0.029 | No |
| fearful-direct | 4 | -0.008 (0.016) | -0.468 (29) | 1 | -0.086 | No |
| fearful-direct | 5 | 0.167 (0.034) | 4.942 (29) | 0.001 | 0.902 | Yes |
| neutral-averted | 1 | -0.004 (0.006) | -0.729 (28) | 1 | -0.136 | No |
| neutral-averted | 2 | 0.003 (0.005) | 0.512 (29) | 1 | 0.094 | No |
| neutral-averted | 3 | -0.015 (0.007) | -2.01 (29) | 1 | -0.367 | No |

| Condition | ms | M (SE) | t(df) | *p*Holm | dz | Sig. |
|---|---|---|---|---|---|---|
| neutral-averted | 4 | 0.005 (0.018) | 0.309 (29) | 1 | 0.056 | No |
| neutral-averted | 5 | 0.061 (0.015) | 3.982 (29) | 0.016 | 0.727 | Yes |
| neutral-direct | 1 | 0.007 (0.003) | 2.059 (28) | 1 | 0.382 | No |
| neutral-direct | 2 | 0.002 (0.006) | 0.413 (29) | 1 | 0.075 | No |
| neutral-direct | 3 | -0.016 (0.008) | -1.92 (29) | 1 | -0.35 | No |
| neutral-direct | 4 | 0.001 (0.018) | 0.07 (29) | 1 | 0.013 | No |
| neutral-direct | 5 | 0.137 (0.025) | 5.551 (29) | < 0.001 | 1.014 | Yes |

*Note.* PAS co-information values are bias-corrected and expressed in bits. Positive values indicate that PAS ratings accounted for shared stimulus-response information; negative values indicate that conditioning on PAS revealed additional stimulus-response dependency. $p_{\text{Holm}}$ = Holm-corrected $p$ value.

### EXPERIMENT 2

#### Supplementary Note 6. Type-1 signal detection sensitivity

The full omnibus type-1 sensitivity analyses are reported in **Supplementary Table S14**. Localisation sensitivity showed strong effects of exposure duration and spatial frequency, as well as a reliable direct-gaze advantage. The gaze effect depended on spatial frequency and exposure duration, consistent with the main-text conclusion that low-spatial-frequency information supported the earliest eye-contact effect. Expression sensitivity showed a main effect of gaze direction, whereas gaze-direction sensitivity showed limited evidence of reliable categorisation under the ultra-brief, spatial-frequency-filtered conditions.

##### Supplementary Table S14

*Omnibus type-1 sensitivity effects in Experiment 2.*

| Task | Effect | F | df | *p* | $\eta_p^2$ |
|---|---|---|---|---|---|
| Location | Expression | 11.79 | 1, 29 | = .002 | 0.29 |
| Location | Gaze direction | 57.82 | 1, 29 | < .001 | 0.67 |
| Location | Spatial frequency | 232.64 | 1, 29 | < .001 | 0.89 |
| Location | Exposure | 538.82 | 1, 29 | < .001 | 0.95 |
| Location | Expression × Gaze direction | 0.05 | 1, 29 | = .827 | 0.00 |
| Location | Expression × Spatial frequency | 1.05 | 1, 29 | = .313 | 0.04 |

| **Task** | **Effect** | **F** | **df** | $p$ | $\eta_p^2$ |
|---|---|---|---|---|---|
| Location | Expression × Exposure | 3.05 | 1, 29 | = .091 | 0.10 |
| Location | Gaze direction × Spatial frequency | 13.97 | 1, 29 | < .001 | 0.33 |
| Location | Gaze direction × Exposure | 5.06 | 1, 29 | = .032 | 0.15 |
| Location | Spatial frequency × Exposure | 0.00 | 1, 29 | = .945 | 0.00 |
| Location | Expression × Gaze direction × Spatial frequency | 0.06 | 1, 29 | = .802 | 0.00 |
| Location | Expression × Gaze direction × Exposure | 0.01 | 1, 29 | = .919 | 0.00 |
| Location | Expression × Spatial frequency × Exposure | 0.42 | 1, 29 | = .520 | 0.01 |
| Location | Gaze direction × Spatial frequency × Exposure | 0.59 | 1, 29 | = .450 | 0.02 |
| Location | Expression × Gaze direction × Spatial frequency × Exposure | 0.84 | 1, 29 | = .368 | 0.03 |
| Expression | Gaze direction | 4.27 | 1, 29 | = .048 | 0.13 |
| Expression | Spatial frequency | 2.83 | 1, 29 | = .103 | 0.09 |
| Expression | Exposure | 0.62 | 1, 29 | = .436 | 0.02 |
| Expression | Gaze direction × Spatial frequency | 1.59 | 1, 29 | = .217 | 0.05 |
| Expression | Gaze direction × Exposure | 2.11 | 1, 29 | = .157 | 0.07 |
| Expression | Spatial frequency × Exposure | 0.05 | 1, 29 | = .817 | 0.00 |
| Expression | Gaze direction × Spatial frequency × Exposure | 1.35 | 1, 29 | = .254 | 0.04 |
| Gaze direction | Expression | 0.81 | 1, 29 | = .376 | 0.03 |
| Gaze direction | Spatial frequency | 1.21 | 1, 29 | = .279 | 0.04 |
| Gaze direction | Exposure | 0.18 | 1, 29 | = .678 | 0.01 |
| Gaze direction | Expression × Spatial frequency | 0.25 | 1, 29 | = .618 | 0.01 |

| Task | Effect | F | df | *p* | $\eta_p^2$ |
|---|---|---|---|---|---|
| Gaze direction | Expression × Exposure | 4.64 | 1, 29 | = .040 | 0.14 |
| Gaze direction | Spatial frequency × Exposure | 0.23 | 1, 29 | = .635 | 0.01 |
| Gaze direction | Expression × Spatial frequency × Exposure | 0.02 | 1, 29 | = .876 | 0.00 |

*Note.* Rows report repeated-measures ANOVA effects for type-1 sensitivity ($d\prime$). SF = spatial frequency. p values are Greenhouse-Geisser corrected where applicable. $\eta_p^2$ = partial eta squared.

Condition-wise tests against zero are reported in **Supplementary Table S15**. Localisation sensitivity was reliable for all low-spatial-frequency conditions at both exposure durations. In the high-spatial-frequency condition, localisation sensitivity was weak or absent at 3 ms for neutral faces but reliable for all conditions at 5 ms. Expression sensitivity was reliable only for direct-gaze low-spatial-frequency faces, whereas gaze-direction sensitivity survived correction only for fearful high-spatial-frequency faces at 5 ms.

##### Supplementary Table S15

*Condition-wise type-1 sensitivity tests against zero in Experiment 2.*

| Task | Condition | SF | Exp. (ms) | M | t(29) | *p*Holm | dz |
|---|---|---|---|---|---|---|---|
| Location | fearful–averted | LSF | 3 | 0.802 | 17.17 | < .001 | 3.13 |
| Location | fearful–direct | LSF | 3 | 0.949 | 14.80 | < .001 | 2.70 |
| Location | neutral–averted | LSF | 3 | 0.692 | 15.15 | < .001 | 2.77 |
| Location | neutral–direct | LSF | 3 | 0.858 | 12.83 | < .001 | 2.34 |
| Location | fearful–averted | LSF | 5 | 1.596 | 30.26 | < .001 | 5.52 |
| Location | fearful–direct | LSF | 5 | 1.817 | 30.85 | < .001 | 5.63 |
| Location | neutral–averted | LSF | 5 | 1.531 | 25.55 | < .001 | 4.66 |
| Location | neutral–direct | LSF | 5 | 1.727 | 25.79 | < .001 | 4.71 |
| Location | fearful–averted | HSF | 3 | 0.095 | 3.14 | = .002 | 0.57 |
| Location | fearful–direct | HSF | 3 | 0.117 | 2.79 | = .005 | 0.51 |
| Location | neutral–averted | HSF | 3 | 0.010 | 0.32 | = .374 | 0.06 |
| Location | neutral–direct | HSF | 3 | 0.023 | 0.65 | = .259 | 0.12 |

| Task | Condition | SF | Exp. (ms) | M | t(29) | *p*Holm | dz |
|---|---|---|---|---|---|---|---|
| Location | fearful–averted | HSF | 5 | 0.865 | 16.02 | < .001 | 2.93 |
| Location | fearful–direct | HSF | 5 | 0.953 | 15.50 | < .001 | 2.83 |
| Location | neutral–averted | HSF | 5 | 0.820 | 11.96 | < .001 | 2.18 |
| Location | neutral–direct | HSF | 5 | 0.957 | 12.88 | < .001 | 2.35 |
| Expression | averted | LSF | 3 | 0.085 | 1.15 | = .259 | 0.21 |
| Expression | direct | LSF | 3 | 0.169 | 2.09 | = .023 | 0.38 |
| Expression | averted | LSF | 5 | 0.027 | 0.30 | = .383 | 0.05 |
| Expression | direct | LSF | 5 | 0.325 | 4.33 | < .001 | 0.79 |
| Expression | averted | HSF | 3 | -0.011 | -0.17 | = .947 | -0.03 |
| Expression | direct | HSF | 3 | 0.028 | 0.37 | = .356 | 0.07 |
| Expression | averted | HSF | 5 | 0.005 | 0.07 | = .947 | 0.01 |
| Expression | direct | HSF | 5 | 0.064 | 0.97 | = .342 | 0.18 |
| Gaze direction | fearful | LSF | 3 | 0.055 | 0.77 | = .222 | 0.14 |
| Gaze direction | neutral | LSF | 3 | 0.031 | 0.47 | = .639 | 0.09 |
| Gaze direction | fearful | LSF | 5 | 0.119 | 1.74 | = .093 | 0.32 |
| Gaze direction | neutral | LSF | 5 | -0.033 | -0.43 | = .666 | -0.08 |
| Gaze direction | fearful | HSF | 3 | 0.078 | 1.32 | = .098 | 0.24 |
| Gaze direction | neutral | HSF | 3 | 0.096 | 1.37 | = .182 | 0.25 |
| Gaze direction | fearful | HSF | 5 | 0.159 | 3.02 | = .005 | 0.55 |
| Gaze direction | neutral | HSF | 5 | 0.072 | 0.88 | = .194 | 0.16 |

*Note.* Rows report one-sample tests against zero for type-1 sensitivity ($d\prime$), Holm-corrected within the relevant family of tests. SF = spatial frequency; Exp. = exposure duration; dz = Cohen's dz.

The direct-minus-averted localisation contrasts are reported in **Supplementary Table S16**. These contrasts show that the direct-gaze advantage was reliable in the low-spatial-frequency condition for both expressions and exposure durations, but emerged in the high-spatial-frequency condition only at 5 ms.

##### Supplementary Table S16

*Direct-minus-averted localisation sensitivity contrasts in Experiment 2.*

| Expression | SF | Exp. (ms) | Mdiff | t(29) | *p*Holm | dz |
|---|---|---|---|---|---|---|
| Fearful | LSF | 3 | 0.147 | 2.93 | = .003 | 0.53 |
| Neutral | LSF | 3 | 0.166 | 2.72 | = .005 | 0.50 |

| **Expression** | **SF** | **Exp. (ms)** | **Mdiff** | **t(29)** | ***p*Holm** | **dz** |
|---|---|---|---|---|---|---|
| Fearful | LSF | 5 | 0.221 | 3.69 | $< .001$ | 0.67 |
| Neutral | LSF | 5 | 0.196 | 3.73 | $< .001$ | 0.68 |
| Fearful | HSF | 3 | 0.022 | 0.50 | $= .309$ | 0.09 |
| Neutral | HSF | 3 | 0.013 | 0.42 | $= .337$ | 0.08 |
| Fearful | HSF | 5 | 0.088 | 2.12 | $= .043$ | 0.39 |
| Neutral | HSF | 5 | 0.138 | 2.93 | $= .006$ | 0.54 |

*Note.* Rows report one-sample tests against zero for the direct-minus-averted difference in localisation $d'$. SF = spatial frequency; Exp. = exposure duration; Mdiff = mean direct-minus-averted difference; dz = Cohen's dz. p values are Holm-corrected.

#### Supplementary Note 7. Response-bias analyses

Response-bias analyses were conducted to assess whether the sensitivity effects could be explained by shifts in response criterion. No omnibus effect reached significance for localisation bias. Significant effects in expression and gaze-direction response bias are summarised in **Supplementary Table S17**. These effects did not mirror the localisation sensitivity pattern and therefore do not account for the early direct-gaze localisation advantage.

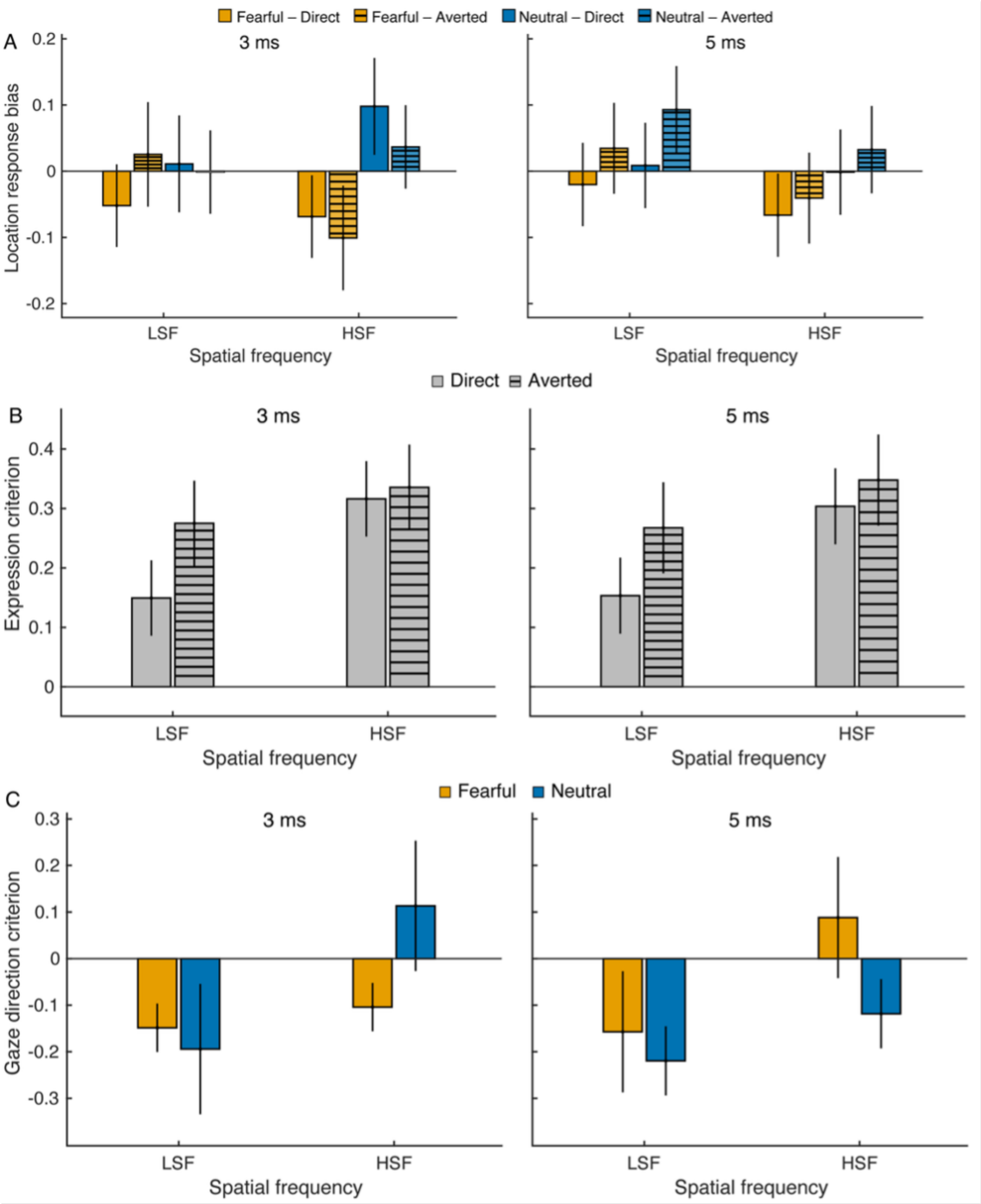


**Supplementary Fig. 5. Response bias and criterion analyses in Experiment 2.** (**A**) Localisation response bias at 3 and 5 ms, plotted by spatial frequency and stimulus condition. (**B**) Expression criterion at 3 and 5 ms, plotted by spatial frequency and gaze direction. (**C**) Gaze-direction criterion at 3 and 5 ms, plotted by spatial frequency and emotional expression. LSF, low-spatial-frequency faces; HSF, high-spatial-frequency faces. Values near zero indicate little systematic response bias or a neutral response criterion. Error bars indicate within-subject SEM. These analyses complement the Experiment 2 sensitivity results by showing that the spatial-frequency-dependent effects were not primarily explained by response-bias or criterion shifts.

#### Supplementary Table S17

*Significant response-bias effects in Experiment 2.*

| Task | Effect | F | df | $p$ | $\eta_p^2$ |
|---|---|---|---|---|---|
| Expression | Gaze direction × Exposure | 9.26 | 1, 29 | = .005 | 0.24 |
| Expression | Spatial frequency × Exposure | 15.20 | 1, 29 | < .001 | 0.34 |
| Gaze direction | Expression | 4.40 | 1, 29 | = .045 | 0.13 |
| Gaze direction | Spatial frequency × Exposure | 4.27 | 1, 29 | = .048 | 0.13 |

*Note.* Only significant omnibus effects are shown to keep the supplementary table focused on interpretable response-bias results. No localisation-bias effect reached significance. $\eta_p^2$ = partial eta squared.

#### Supplementary Note 8. Information-theoretic localisation analyses

Localisation mutual-information tests are reported in **Supplementary Table S18**. Consistent with the type-1 signal detection analyses, localisation responses carried reliable stimulus-location information for low-spatial-frequency faces at both 3 ms and 5 ms. For high-spatial-frequency faces, localisation information did not exceed the permutation-corrected baseline at 3 ms but became reliable at 5 ms.

##### Supplementary Table S18

*Condition-wise localisation mutual information in Experiment 2.*

| Expression | Gaze direction | SF | Exp. (ms) | MI (bits) | t(29) | $p$Holm | dz |
|---|---|---|---|---|---|---|---|
| Fearful | Averted | LSF | 3 | 0.109 | 7.33 | < .001 | 1.34 |
| Fearful | Direct | LSF | 3 | 0.165 | 7.59 | < .001 | 1.39 |
| Neutral | Averted | LSF | 3 | 0.079 | 6.38 | < .001 | 1.17 |
| Neutral | Direct | LSF | 3 | 0.137 | 5.25 | < .001 | 0.96 |
| Fearful | Averted | LSF | 5 | 0.423 | 16.88 | < .001 | 3.08 |
| Fearful | Direct | LSF | 5 | 0.541 | 16.31 | < .001 | 2.98 |
| Neutral | Averted | LSF | 5 | 0.394 | 14.10 | < .001 | 2.57 |
| Neutral | Direct | LSF | 5 | 0.491 | 14.03 | < .001 | 2.56 |
| Fearful | Averted | HSF | 3 | -0.028 | -15.16 | = 1.000 | -2.77 |
| Fearful | Direct | HSF | 3 | -0.019 | -6.52 | = 1.000 | -1.19 |
| Neutral | Averted | HSF | 3 | -0.027 | -14.42 | = 1.000 | -2.63 |
| Neutral | Direct | HSF | 3 | -0.027 | -14.04 | = 1.000 | -2.56 |
| Fearful | Averted | HSF | 5 | 0.135 | 6.55 | < .001 | 1.20 |
| Fearful | Direct | HSF | 5 | 0.159 | 6.20 | < .001 | 1.13 |
| Neutral | Averted | HSF | 5 | 0.123 | 4.46 | = .001 | 0.82 |
| Neutral | Direct | HSF | 5 | 0.169 | 4.89 | < .001 | 0.89 |

*Note.* Rows report one-sample tests against zero for bias-corrected mutual information between stimulus location and localisation response. Negative values can occur after permutation-based bias correction and indicate no evidence for information above the null baseline. SF = spatial frequency; Exp. = exposure duration; MI = mutual information; dz = Cohen's dz. p values are Holm-corrected.

Direct-minus-averted mutual-information contrasts are reported in **Supplementary Table S19**. Direct gaze increased localisation information most clearly in the low-spatial-frequency condition, with reliable effects for fearful faces at 3 ms and for both fearful and neutral faces at 5 ms. None of the high-spatial-frequency direct-minus-averted contrasts survived correction.

**Supplementary Table S19**

*Direct-minus-averted localisation mutual-information contrasts in Experiment 2.*

| **Expression** | **SF** | **Exp. (ms)** | **ΔMI (bits)** | **t(29)** | ***p*Holm** | **dz** |
|---|---|---|---|---|---|---|
| Fearful | LSF | 3 | 0.056 | 3.12 | = .028 | 0.57 |
| Neutral | LSF | 3 | 0.058 | 2.43 | = .130 | 0.44 |
| Fearful | LSF | 5 | 0.118 | 3.61 | = .009 | 0.66 |
| Neutral | LSF | 5 | 0.097 | 3.55 | = .010 | 0.65 |
| Fearful | HSF | 3 | 0.008 | 2.26 | = .175 | 0.41 |
| Neutral | HSF | 3 | 0.001 | 0.34 | = 1.000 | 0.06 |
| Fearful | HSF | 5 | 0.024 | 1.42 | = .829 | 0.26 |
| Neutral | HSF | 5 | 0.046 | 2.59 | = .096 | 0.47 |

*Note.* Rows report one-sample tests against zero for the direct-minus-averted difference in localisation mutual information. ΔMI = mean direct-minus-averted difference in bias-corrected mutual information; SF = spatial frequency; Exp. = exposure duration; dz = Cohen's dz. *p* values are Holm-corrected.
